# Cell type purification by single-cell transcriptome-trained sorting

**DOI:** 10.1101/502773

**Authors:** Chloé S Baron, Aditya Barve, Mauro J Muraro, Gitanjali Dharmadhikari, Reinier van der Linden, Anna Lyubimova, Eelco J. P. de Koning, Alexander van Oudenaarden

**Author notes:** These authors contributed equally (Chloé S Baron, Aditya Barve, Mauro J Muraro). Correspondence should be addressed to A.v.O.

## Abstract

Traditional cell type enrichment using fluorescence activated cell sorting (FACS) relies on methods that specifically label the cell type of interest. Here we propose GateID, a computational method that combines single-cell transcriptomics for unbiased cell type identification with FACS index sorting to purify cell types of choice. We validate GateID by purifying various cell types from the zebrafish kidney marrow and the human pancreas without resorting to specific antibodies or transgenes.

---

The ability to enrich for different cell types from heterogeneous tissues underpins much of current biological and clinical research. Methods using FACS to enrich cells use reporter transgenes or fluorescent antibodies that are specific for the cell type of interest. However, limited availability of specific antibodies or - in case of reporter constructs - the need for genetic manipulation, limit this approach, especially in the case of human tissues. Here, we describe GateID, an optimization algorithm that combines single-cell FACS and transcriptome information with a goal to predict FACS gates for cell types that were identified by the single-cell mRNA-sequencing (scRNA-Seq). It benefits from two technological breakthroughs: single-cell transcriptomics and FACS index sorting. Recent studies have demonstrated that a combining single-cell transcriptomics with index sorting can be used to better characterize cellular subpopulations^1–3^. Conversely, we reasoned that single-cell transcriptome data could be used to improve sorting of pure cell populations. Given a single-cell transcriptome training dataset GateID predicts novel FACS gates based on general properties such as cell size and granularity, nuclear staining, cellular proliferation, and mitochondrial activity. GateID allows the purification of specific cell types or states without the use of specific markers such as antibodies or transgenes. Importantly, GateID prediction is solely data-driven and does not require *a priori* information about FACS gates or cellular markers.

The GateID workflow starts with generating a training dataset of the organ/tissue of interest (**Fig. 1, step 1**). To this end, single live cells are sorted while recording index data in all available scatter and fluorescent channels (**Fig. 1, step 1a**). Next, the transcriptome of all sorted single cells is sequenced using SORT-Seq^4^ and the cell type composition of the organ/tissue of interest is determined (**Fig. 1, step 1b**). The GateID training dataset is generated by merging the index sorting parameters with the cell type information obtained by scRNA-Seq for each cell (**Fig. 1, step 1c**). After defining the desired cell type, the computational gate design occurs (**Fig. 1, step 2**). At the core of GateID is an optimization algorithm that attempts to predict gates to obtain the maximum number of desired cells while minimizing the number of undesired cells. It iterates this procedure through all combinations of FACS channels and subsequently through combinations of gates to predict best gates in terms of purity and yield (see Online Methods, **Fig. 1, step 2d-e**). Finally, the GateID predicted gates are experimentally validated using a new sample of the organ/tissue of interest (**Fig. 1, step 3**). Predicted gates are normalized to the new experimental dataset to correct for biological inter-individual variability and FACS technical variability (**Fig. 1, step 3f**). Single cells passing through normalized GateID gates are sorted and sequenced using scRNA-Seq (**Fig. 1, step 3g-h**). The cell type composition of the GateID enriched library is determined and the experimental purity of the GateID gates is calculated.

**Figure 1.**
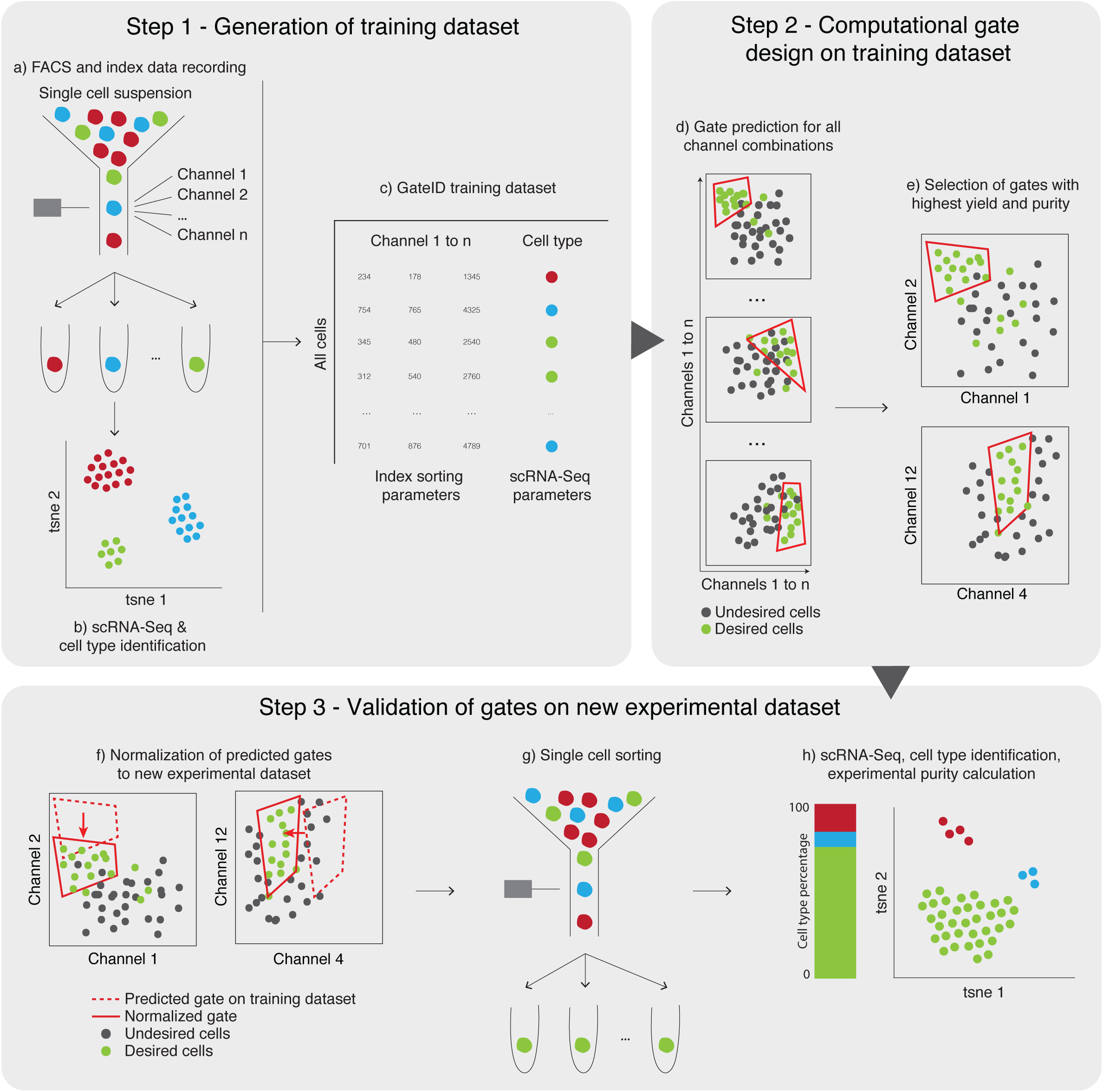
GateID workflow. In step 1, the GateID training dataset is generated. **(a)** Live single cells from the organ of interest are sorted in an unbiased manner and index data for all available channels is recorded. **(b)** Single cells sorted in (a) are sequenced to determine the cell type composition of the organ. **(c)** The GateID training dataset is generated after merging the FACS index data and the cell type information for each single cell. In step 2, the gates are computationally designed for the desired cell type. **(d)** Gates are computed for each possible combination of channels. **(e)** The best combination of gates is chosen in order to maximize yield and purity of the desired cell type. In step 3, GateID predicted gates are experimentally tested. **(f)** The predicted gates are normalized to the new experimental dataset. **(g)** Single cells in GateID gates are sorted. **(h)** After scRNA-Seq, cell types present in the GateID enriched library are determined and the experimental purity is calculated.

To test our method, we first decided to focus on the adult zebrafish whole kidney marrow (WKM), the primary site of production of hematopoietic cells in zebrafish. Traditionally, their isolation relies on a limited number of antibodies, transgenic lines or manual gating subject to high variability^5,6^. To check if GateID could be a more attractive method, we first generated a training dataset of single live WKM hematopoietic cells (DAPI-) by merging FACS index data in 12 dimensions (scatter and fluorescent dimensions from a BD FACSJazz™) and cell type information for 1252 cells from 3 zebrafish. Using cell clustering and known markers, we identified 7 hematopoietic cell types^7–11^ (**Supplementary Fig. 1a-c**, see Online Methods) and first aimed to enrich for eosinophils. GateID predicted a yield of 46.9% and a purity of 79.3% to isolate eosinophils using a combination of two gates (**Fig. 2a, Supplementary table 1**). To experimentally validate the predicted gates, we sorted enriched (GateID) and unenriched (live cells) single cells from three independent WKMs (**Fig. 2b**, representative example for WKM 1 to 3). As described above, GateID predicted gates were normalized to each new WKM in order to correct for inter-individual and technical variability (**Fig. 2b**, black gates are non-normalized while red gates are normalized to WKM 2 experimental dataset, see Online Methods). After scRNA-Seq of enriched and unenriched eosinophil libraries, we aimed to calculate experimental purities. To ensure high confidence in our cell type identification and our purity estimates, we clustered all zebrafish GateID experiments together (WKM 1-15, training datasets 1-3) resulting in 15984 single cells (**Fig. 2c**, see Online Methods). We then calculated the experimental purities of all our WKM experiments based on this full dataset. The above-mentioned eosinophil enrichment experiments achieved an experimental purity between 68.9% and 78% purity even with as low eosinophil content as 0.6% in the unenriched population (**Fig. 2d**, barplots, *n*=3). Two experiments of the three (WKM 1, 78% purity and WKM 3, 74.7% purity) show reasonable alignment with predicted purity. The reason the experimental purity varies from predicted, especially in WKM 2, lies in the individual variation that can be observed in both cell type composition and in FACS measurements per experiment (**Supplementary Fig. 2a**). Interestingly, we observed that contaminating cells intermingled with enriched eosinophils in FACS space and are thus difficult to eliminate (**Supplementary fig. 2b**). The contaminating population in all experiments consisted mainly of monocytes. This is not surprising, since eosinophils and myeloid cells occupy partly overlapping FACS regions^12^. Importantly, the enriched eosinophils from each experiment clustered with the eosinophils in the unenriched population (**Fig. 2d**, black and orange points on t-SNE maps, respectively), meaning that GateID enriched cells capture the existing transcriptional variance in eosinophils from the unenriched library. This shows that GateID does not bias for a subpopulation of eosinophils. Finally, to compare GateID to manual gating, we isolated eosinophils as previously described^12^ (**Supplementary Fig. 2c**). This manual gating yielded lower enrichment compared to GateID and revealed a stronger myeloid contamination (**Supplementary Fig. 2d-e**).

**Figure 2.**
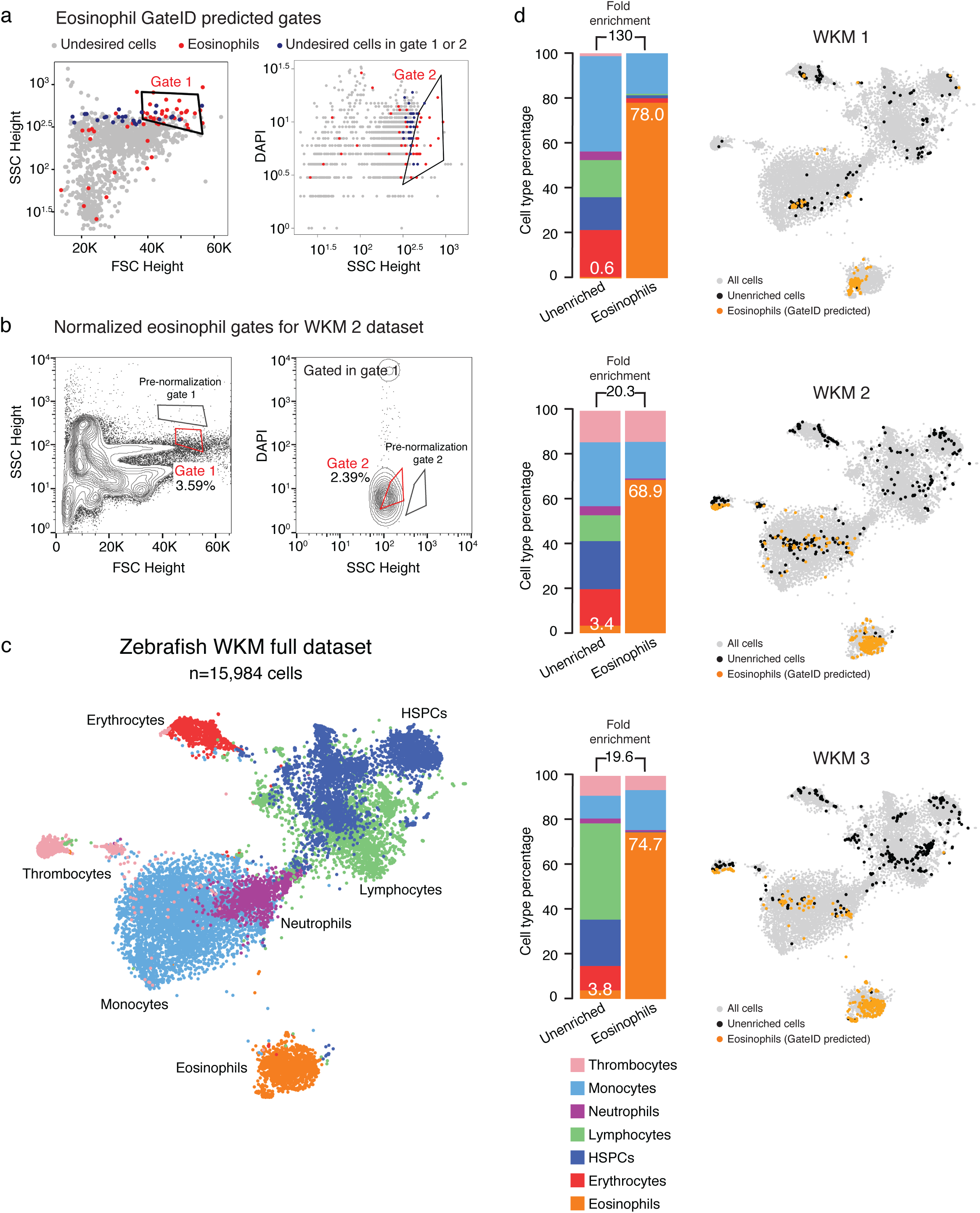
Proof of principle: enrichment of zebrafish eosinophils using GateID. **(a)** GateID predicted gates to isolate eosinophils from unstained WKM on BD FACSJazz™. Gates were predicted on unstained WKM training dataset 1. Red points show desired cells (eosinophils) present in training dataset 1 and blue points show undesired cells present in the other gate. A blue colored undesired cell inside a gate denotes an impure cell that will be sorted. **(b)** Contour plots of unstained WKM cells showing experimental sorting gates for eosinophils for WKM 2 experiment (representative example for WKM 1 to 3 eosinophil enrichment experiments) on BD FACSJazz™. Gates in black represent GateID predicted gates prior to normalization, whereas red gates show GateID normalized sorting gates. Sorted cells passed through normalized gate 1 and gate 2. Percentages of events within each gate are indicated. **(c)** t-SNE map of the complete zebrafish WKM dataset (all WKM training datasets and enrichment experiment datasets of this study, *n*=15.984 cells). Single cells are colored based on cell type. **(d)** Barplots and t-SNE maps showing the outcome of GateID eosinophil enrichments for three independent experiments (WKM 1 to 3) on BD FACSJazz™. Gates were predicted on unstained training dataset 1. In the barplots, numbers within the bars indicate the percentage of eosinophils in the corresponding library and numbers above the bars indicate the cell type fold enrichment between unenriched and GateID enriched library. On the t-SNE maps, grey points represent all cells from the WKM dataset. For each experiment, black dots are single cells in the unenriched library for a given experiment, while colored dots are single cells in the GateID enriched library for the same experiment.

We next aimed to isolate additional hematopoietic cell types from the WKM. Our first BD FACSJazz™ training dataset (**Supplementary Fig. 1b**) was obtained using DAPI^−^ WKM cells with a limited number of cells per individual. While this dataset was sufficient to design gates for eosinophils, GateID was unable to predict gates with satisfying purity and yield for HSPCs, lymphocytes or monocytes. Indeed, plotting purity versus yield for the best combination of GateID gates for each of these cell type clearly showed that training dataset 1 did not allow us to enrich these cell types to satisfying purities (**Supplementary Fig 1d**). We hypothesized that this limitation could be resolved by enhancing cell type separation in FACS space. However, we aimed to keep GateID antibody and transgene-free and therefore chose to stain WKM cells with generic cellular dyes. We chose MitoTracker, a fluorescent die that reflects mitochondrial abundance and activity, and CFSE, which binds to cytoplasmic proteins (see Online Methods). Neither dye stains any one cell type specifically, but all cells. To validate our approach, we used a new WKM sample and split it in two parts, one of which was stained with MitoTracker, CFSE and DAPI, while the other was stained only with DAPI. We FACS sorted and performed scRNA-Seq on both libraries. After identifying the relevant cell types based on the transcriptome, we evaluated all two-gate combinations for the enrichment of HSPCs, lymphocytes, monocytes and eosinophils in MitoTracker+ CFSE+ DAPI^−^ (referred to as “stained” here onwards) and DAPI^−^(referred to as “unstained” here onwards) samples (**Fig. 3a**). We observed that unstained samples allowed to predict gates with lower yield compared to the stained samples. It is important to note that these samples are limited in the number of cells (~250 for each sample) and do not represent possible contaminating cells that could be observed in a larger sample, thereby predicting higher purities than would be obtained experimentally or with a larger dataset.

**Figure 3.**
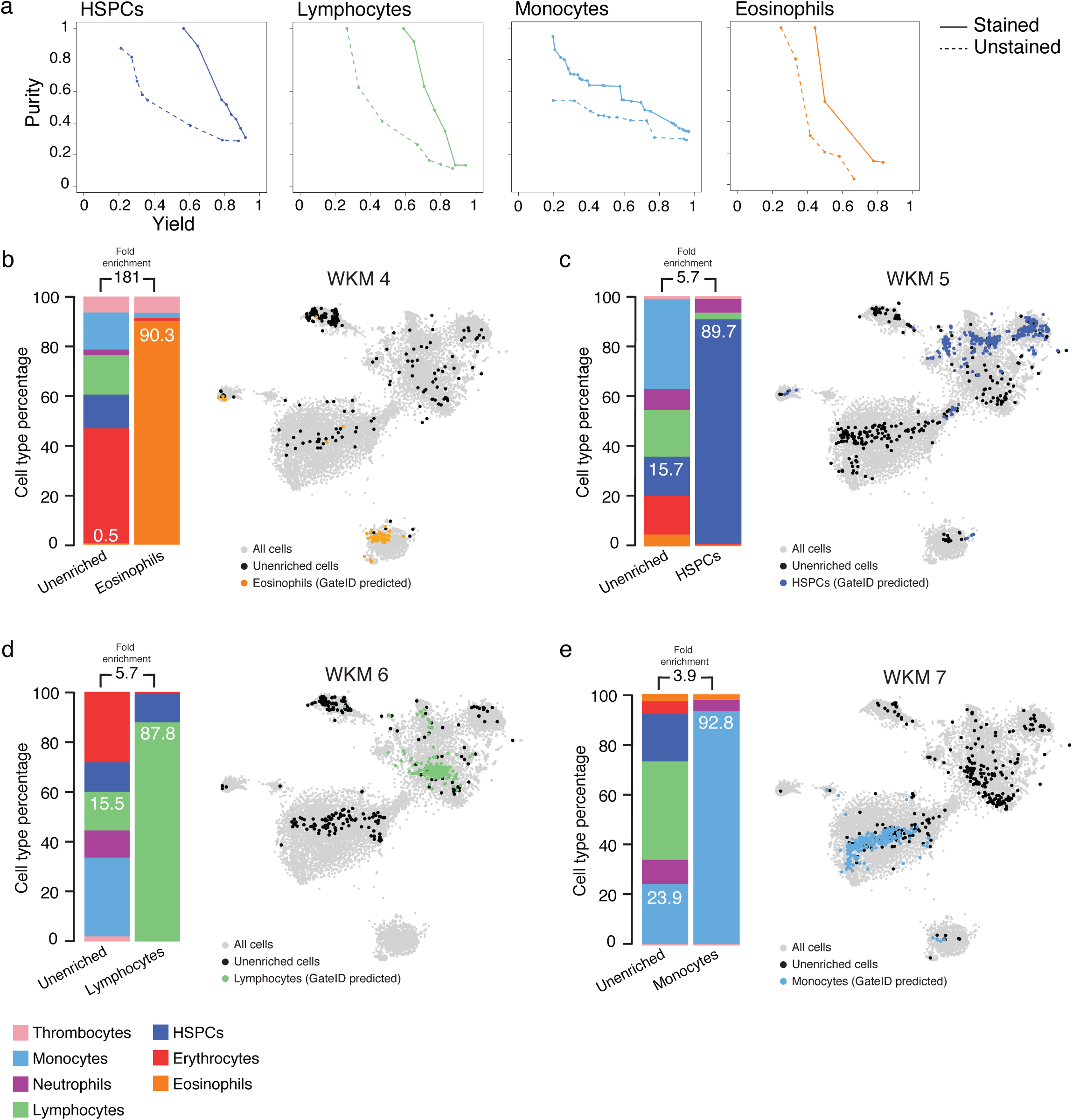
General dyes enhance cell type segregation in FACS space to allow their purification with GateID. **(a)** Curves showing trade-off between yield and purity of GateID solutions for HSPCs, lymphocytes, monocytes and eosinophils on stained (solid line) and unstained (dashed line) cells from the same zebrafish WKM (WKM 7). **(b-e)** Barplots and t-SNE maps showing the outcome of GateID enrichments of **(b)** eosinophil (WKM 4) **(c)** HSPC (WKM 5) **(d)** lymphocyte (WKM 6) and **(e)** monocyte (WKM 7) on BD FACSJazz™. Gates were predicted on stained training dataset 2. In the barplots, numbers in the bars indicate the percentage of the desired cell type in the corresponding library and numbers above the bars indicate the cell type fold enrichment between unenriched and GateID enriched library. On the t-SNE maps, grey points represent all cells from the WKM dataset. For each experiment, black dots are single cells in the unenriched library for a given experiment, while colored dots are single cells in the GateID enriched library for the same experiment.

Since MitoTracker and CFSE allowed better separation of cell types in FACS space, we generated two training datasets of stained WKM hematopoietic cells. WKM training dataset 2 was generated on a BD FACSJazz^TM^ and resulted in 1.201 cells (**Supplementary Fig. 3a**) while training dataset 3 was generated on a BD FACSInflux™ and composed of 1036 cells (**Supplementary Fig. 3b**). The optical setups of our BD FACSJazz™ and BD FACSInflux™ allowed recording of index data in 12 and 27 dimensions, respectively. We used both datasets to design gates to enrich multiple hematopoietic cell types and demonstrate that GateID performance would be independent of the FACS machine of use. First, we repeated the eosinophil enrichments using stained WKM cells and sorting with a BD FACSJazz^TM^ (**Supplementary Fig. 3c-d**). Notably, while GateID predicted gates are unconventional and humanly unintuitive, we show that our enriched population maps back in the same region as the classical manual FACS gate^12^ (**Supplementary Fig. 3d**). We obtained marginally higher purities (85.4% on average) when compared to unstained cells (73.9% on average) (**Fig. 3b, Supplementary Fig. 3e-f**, *n*=3). Next, we used GateID to predict gates to enrich for HSPCs on BD FACSJazz™ and BD FACSInflux™. GateID predicted a yield of 20% and a maximum purity of 90.5% to isolate HSPCs on a BD FACSJazz™ using a combination of two gates, one of them using the MitoTracker fluorescent channel (**Supplementary Fig. 4a**). Not surprisingly, the projection of GateID enriched HSPCs on the classical dimensions of FSC height and SSC height is similar to what is published^12^ (**Supplementary Fig. 4b**). Experimentally, we were able to enrich HSPCs to an average purity of 89% wherein enriched HSPCs clustered together with the unenriched HSPC population for each experiment when visualized on t-SNE (**Fig. 3c, Supplementary Fig. 4c-d**, *n*=3). Additionally, GateID predicted a yield of 30% and purity of 98.6% to isolate HSPCs on a BD FACSInflux™ (**Supplementary Fig. 5a-b**). We obtained purities averaging 67% and observed no bias towards a subset of HSPCs upon GateID enrichment (**Supplementary Fig. 5c-e**, *n*=3). Importantly, we compared our transcriptomics method for cell type calling to manual histological classification to calculate purities. We found high correlation between both methods to calculate HSPC purities after enrichment using GateID (**Supplementary Fig. 5f**, dark blue points for HSPC enrichments on BD FACSJazz™ (triangles) and BD FACSInflux™ (circles)). Finally, to benchmark GateID, we compared it to a classical method of enriching HSPCs based on their low expression of cd41 (**Supplementary Fig. 4e**)^13,14^. Enriched HSPCs from the cd41^low^ fraction from cd41-EGFP transgenic zebrafish yielded an inferior purity compared to GateID predicted gates (**Supplementary Fig. 4f-g**). Surprisingly, the enriched HSPCs were contaminated by neutrophils. This result suggested that neutrophils reside partially in the cd41^low^ WKM fraction, an observation that would have gone undetected without the combination of single-cell FACS and transcriptome information. Next, we used GateID to isolate lymphocytes, using four FACS dimensions including the CFSE fluorescent channel (**Supplementary Fig. 6a-b**). Experimentally, with BD FACSJazz™, we obtained unbiased enrichment between 77% and 91.7% (Fig. 3d, **Supplementary Fig. 6c-e**, *n*=4). *In silico*, we tested the efficiency of lymphocyte manual gating as lymphocytes are characterized by their small FSC height and SSC height properties^12^ (**Supplementary Fig. 6f**). The manual gate yielded 60.9% purity and exhibited HSPC contamination (**Supplementary Fig. 6g**). We then challenged GateID to isolate a subset of myeloid cells on both BD FACSJazz™ and BD FACSInflux™. Neutrophils and monocytes are strongly intermingled in side scatter height vs. forward scatter height^12^. However, GateID made use of the CFSE or MitoTracker dimensions to design gates to purify monocytes (**Supplementary Fig. 7a-b** for BD FACSJazz™ and **Supplementary Fig. 8a-b** for BD FACSInflux™). We succeeded in enriching monocytes to average purities of 79.7% on BD FACSJazz™ and 87.1% on BD FACSInflux™ (**Fig. 3e, Supplementary Fig. 7c-d** for BD FACSJazz™, *n*=3 and **Supplementary Fig. 8c-e** for BD FACSInflux™, *n*=3). We find the enriched populations to overlap with the one present in the live population in t-SNE space for all experiments and found neutrophils to be the highest source of contamination. Finally, as described above, monocyte purities determined by histological classification highly correlated with the one determined by scRNA-Seq (**Supplementary Fig. 5f**, light blue circles). Overall, we demonstrate GateID’s ability to enrich multiple zebrafish hematopoietic cells to high purity solely using generic dyes. GateID proved more robust for cell types we enrich here when compared to the tested manual gating strategies that use FACS scatter properties or fluorescent transgenic lines. In addition, we show that GateID can be successfully used with FACS machines with different optical setups.

Additionally, we used GateID in a human, clinical setting. We and others previously sequenced single cells from islets of Langerhans obtained from human cadaveric material (reviewed in ^15^) to describe the transcriptomes of the 6 major pancreatic cell types (alpha, beta, delta, PP, acinar and ductal cells) implicated in the pathogenesis of diabetes. Unfortunately, isolating live alpha, beta and delta cells to high purity remains a challenge due to the absence of reliable markers. Previous efforts^16^ in obtaining enriched populations of alpha and beta cells by using antibodies are unclear, as delta cell markers were found in the enriched beta cell population, indicating a strong contamination from delta cells (table 1 in ^16^). We thus set out to use GateID to enrich alpha and beta cells to high purity from human pancreas. First, we used one of the donors from our previous dataset^4^ (D30) as a GateID training dataset. We merged the BD FACSJazz™ index parameters to the cell type information for 664 DAPI stained single cells (**Supplementary Fig. 9a**, pancreas training dataset 1). GateID predicted 43% yield and 100% purity for alpha cells and 52% yield and 100% purity for beta cells (**Supplementary Fig. 9b,d**). To experimentally validate the GateID predicted gates, we used a new donor (donor 1) to sort enriched (GateID) and unenriched cells (**Supplementary Fig. 9c,e**). As we did for the zebrafish WKM, we clustered all the scRNA-Seq data from our pancreas experiments to confidently call cell types and calculate the GateID experimental purities. This combined dataset resulted in 10176 cells representing 8 distinct pancreatic cell types (**Fig. 4a**). For donor 1, we obtained 100% pure alpha cell population and a 79% pure beta cell population (**Fig. 4b**, barplot). Importantly, the GateID enriched alpha and beta cells clustered together with the unenriched population, revealing unbiased enrichment of both cell types (**Fig. 4b**, t-SNE maps). We observed that the contamination in donor 1 beta cell gates stemmed from all other pancreatic cell types indicating an overall inefficient exclusion of undesired cell types. We hypothesized that training dataset 1 did not contain enough information about the undesired cells that would be present in an experimental sort or a larger dataset. Thus, GateID would predict gates without that specific information and such gates may not exclude undesired cells efficiently. To test this hypothesis, we built a larger training dataset of 2255 cells by sorting DAPI stained single cells on a BD FACSJazz^TM^ and performing scRNA-Seq to identify the main pancreatic cell types (**Supplementary Fig. 10a**, pancreas training dataset 2). First, we repeated the alpha cell enrichment with new predicted gates designed on training dataset 2 (yield 51% and purity 97%, **Supplementary Fig. 10b-c**) and obtained 89% experimental purity (**Supplementary Fig. 10g**). Next, GateID predicted gates of 26% yield and 98% purity for beta cells (**Supplementary Fig. 10d**). We experimentally validated these gates with three independent donors (donors 2–4) and achieved an average purity of 95% (Fig. 4c, **Supplementary Fig. 10e-g**). In t-SNE space, GateID enriched beta cells did not separate from the ones in the unenriched fraction. This means that GateID enriched beta cells capture the different transcriptional profiles present in the unenriched sample revealing unbiased enrichment of beta cells. Overall, our results show that GateID can faithfully predict gates to enrich for alpha and beta cells from the human pancreas, allowing us to purify these cell types solely based on their intrinsic FACS scatter and autofluorescence properties.

**Figure 4.**
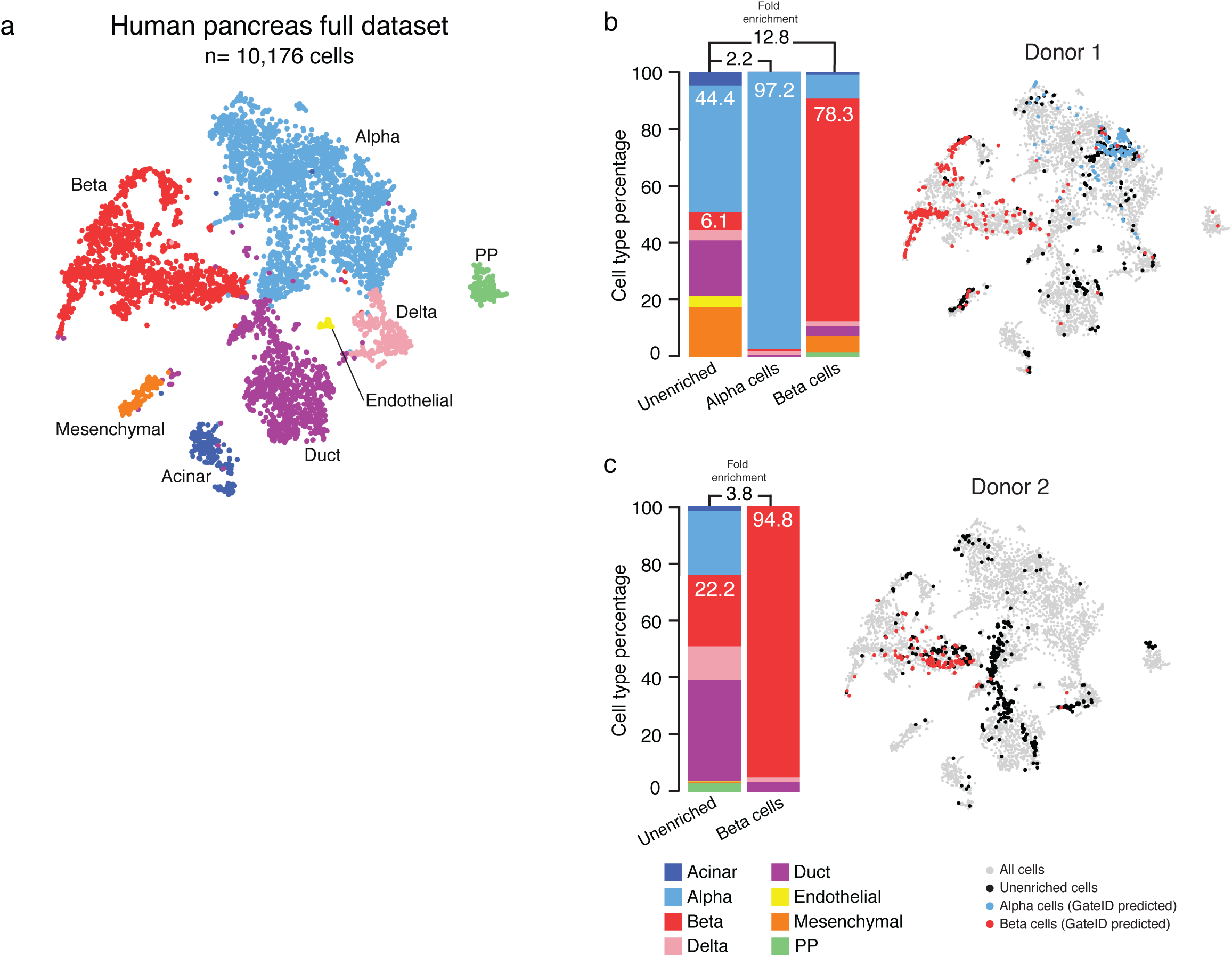
GateID allows enrichment of α and β cells from human pancreatic islets. **(a)** t-SNE map of the complete pancreas dataset (all pancreas training datasets and enrichment experiment datasets, *n*=10.176 cells). Single cells are colored based on cell type. **(b-c)** Barplots and t-SNE maps showing the outcome of GateID alpha and beta cell enrichments for two independent donors on BD FACSJazz™. Gates for donor 1 were predicted on unstained training dataset 1 and gates for donor 2 were predicted on unstained training dataset 2. In the barplots, numbers within the bars indicate the percentage of alpha and beta cells in the corresponding library and numbers above the bars indicate the cell type fold enrichment between unenriched and GateID enriched library. On the t-SNE maps, grey points represent all cells from the pancreas dataset. For each experiment, black dots are single cells in the unenriched library for a given experiment, while colored dots are single cells in the GateID enriched library for the same experiment.

In short, we have described a novel computational method that combines single-cell transcriptomics and single-cell FACS to predict FACS gates that allow cell type enrichment without the aid of transgenes or antibodies. To demonstrate the effectiveness of GateID, we enriched four major hematopoietic cell types from the zebrafish WKM, a tissue for which transgenes labeling specific cell types are labor intensive to generate and antibodies are limited. Our approach proves sufficiently robust to enrich for hematopoietic cell types ranging from 0.5% (eosinophils) to 35% (monocytes) of the total WKM cell composition (Fig. 2–3). Importantly, we show that the performance of GateID does not depend on the proportion of the desired cell type in the tissue of interest. Indeed, no correlation (Pearson’s r = 0.07) was found between the achieved experimental purity and the abundance of the cell type of interest in all our 22 WKM enrichment experiments (**Supplementary Fig. 11a**). The performance of GateID is also independent of age and gender, as we did not control for these parameters while choosing our zebrafish or human samples. Additionally, our approach allows purification of more than one cell type from one animal as shown by purifying eosinophils, lymphocytes and monocytes from WKM 8 (**Supplementary Fig. 3e, 6d, 7d**). Finally, GateID performs better than classical methods of cell type enrichment as shown for examples concerning eosinophils, HSPCs and lymphocytes (**Supplementary Fig. 2d-e, 4e-g, 6f-g**).

In the WKM, we demonstrate that the distinction of the desired cell type is in FACS space compared to other cell types influenced the performance of GateID. A limitation of GateID is that it may not always perform well if cell types largely overlap in FACS space. That is, in absence of a specific marker for the cell type of choice (e.g. antibody or fluorescent reporter), cell types might be difficult to segregate based on their scatter and autofluorescence properties alone. While these properties were sufficient to predict GateID gates for zebrafish eosinophils, they were not sufficient for zebrafish lymphocytes, HSPCs and monocytes (**Supplementary Fig. 1d**). While remaining antibody and transgene-free, we demonstrated that general dyes (MitoTracker and CFSE) allow segregation of hematopoietic cell types in FACS space and allow successful gate prediction and validation (**Fig. 2**). Finally, we want to emphasize that GateID can be used in combination with any antibody or transgene of choice. This approach would allow refinement of manual gating and potentially intricate gating for subsets of cells without the need for large antibody panels.

Additionally, we showed that GateID could also enrich human alpha and beta cells from the islets of Langerhans. This is especially important for human tissues where purification of cell types is completely restricted to the availability of antibodies. Here, we demonstrate that GateID allows to purify live alpha and beta cells in an antibody independent manner. In our pancreas enrichment experiments, we demonstrated the role of biological variability between training dataset and all experimental datasets and its role on GateID’s performance. Variability mainly springs from variable proportion of cell types and different statistical properties for each cell type in different datasets. GateID offers a normalization strategy and we showed good robustness to correct for such variability. However, GateID gate prediction and normalization will perform adequately only if the training dataset captures sufficient diversity from the chosen sample. Along these lines, we observed that insufficient knowledge of the FACS properties of contaminating cells jeopardized GateID’s performance to enrich for beta cells from pancreas training dataset 1 (**Fig. 4b**). By increasing the size of this training dataset, GateID could more efficiently gate out contaminating cells, which was reflected in an increase in experimental purities to 99.3% (**Fig. 4c**, **Supplementary Fig. 10f,g**). To more precisely estimate the adequate size of a training dataset, we performed beta cell gate design using GateID on various datasets computationally generated from pancreas training dataset 1 (**Supplementary Fig. 11b**, see online method). Importantly, we computationally changed the ratio of contaminating cells in the enlarged datasets to visualize the impact of the proportion of non-beta cells on the performance of GateID gates. We observed that gates designed on a smaller dataset (1x training dataset 1, 664 cells) fare poorly in comparison to gates designed on a larger dataset (2x and 3x training dataset 1, 1328 and 1992 cells respectively). Specifically, increasing the training dataset two fold to 1356 cells ensures higher mean purity even in the case of twice the amount of contaminating cells and may ensure higher robustness to fluctuations in cell proportions. Overall, in line with our results with WKM training datasets, we find training datasets ranging from 1000 to 1300 to allow robust gate design using GateID. Whereas generating such a training dataset can be a limitation due to the costs of scRNA-Seq, we emphasize that training datasets can be used to generate gates for all cell type present in the organ of choice. Additionally, thanks to GateID robust normalization strategy, the user will be able to enrich for a desired cell type in an unlimited amount of experiments, on different samples.

Overall, we envisage a broad application of GateID to make purification of any given cell type easier and to allow enrichment of cell types never isolated before.

## Methods

### Ethical statement

All animal experiments were performed in accordance with institutional and governmental regulations, and were approved by the Dier Experimenten Commissie of the Royal Netherlands Academy of Arts and Science and performed according to the guidelines.

Human cadaveric donor pancreata were procured through a multi-organ donor program. Pancreatic tissue was only used if the pancreas could not be used for clinical pancreas or islet transplantation, only if research consent was given and according to national laws.

### Tissue isolation

The WKM of WT and cd41-GFP zebrafish were isolated as described previously^1^. Briefly, after a ventral midline incision the internal organs were removed. The kidney was carefully dissected and collected in PBS supplemented with FCS. To mechanically dissociate single hematopoietic cells, the tissue was passed multiple times through a 1 ml low-bind pipet tip. The cells were filtered (70um and 40um cell strainers (VWR)) and washed. The pellet of hematopoietic cells was resuspended in PBS/FCS supplemented with DAPI (dilution 1/2000, Thermo Fisher) to assess cell viability. In case of staining, the pellet of hematopoietic cells was resuspended in PBS/FCS supplemented with both MitoTracker and CFSE (dilution 1/4000) and incubated at room temperature for 10 minutes. Cells were washed and resuspended in PBS/FCS supplemented with DAPI as described above. For training dataset generation, DAPl^−^ single cells were sorted (BD FACSJazz™ or BD FACSlnflux™) and erythrocytes with low forward and side scatter were excluded as described in Supplementary Fig. 1a. For GateID enrichment experiments, cells passing through all gates were sorted. For histology, pools of 10.000 cells were sorted in PBS supplemented with FCS and fixed 10 minutes in 4% PFA. After washing, cytospins were performed as described in ref. 5. Cells were post-fixed on slide and May-Grunwald-Giemsa staining was performed following manufacturer’s instructions. Human pancreas isolation was done as described previously^2^.

### Single-Cell mRNA Sequencing of Single Cells

We used SORT-seq^2^ to sequence the transcriptome from single cells and store FACS information from single cells (index files). All sorts were carried out using BD FACSJazz^TM^ or BD FACSlnflux™. Unless mentioned otherwise, we used the following protocol for both model systems mentioned in this study. We lysed cells by incubating them at 65°C for 5 minutes, and then used Nanodrop II liquid handling platform (GC biotech) to dispense RT and second strand mixes. The aqueous phase was separated from the oil phase after pooling all cells into one library, followed by IVT transcription. The CEL-Seq2 protocol was used for library prep^3^. Primers consisted of a 24 bp polyT stretch, a 4 or 6bp random molecular barcode (UMI), a cell-specific 8bp barcode, the 5’ Illumina TruSeq small RNA kit adaptor and a T7 promoter. We used TruSeq small RNA primers (Illumina) for preparation of Illumina sequencing libraries and then paired-end sequenced them at 75 bp read length using Illumina NextSeq at approximately 45 million and 30 million reads for zebrafish kidney marrow and human pancreatic libraries respectively.

### Data analysis

Zebrafish WKM and human pancreas were analyzed separately as follows. For each model system we analyzed, paired-end reads were aligned to the transcriptome of that model system using BWA^5^. We used Read 1 for assigning reads to correct cells and libraries, while read 2 was mapped to gene models. Only reads mapping to unique locations were kept. We corrected read counts for UMI barcodes by removing duplicate reads that had identical combinations of library, cellular, and molecular barcodes and were mapped to the same gene. Transcripts were counted using 256 UMI barcodes for the human pancreas (donor 1) and 4096 UMI barcodes for the other human donors and the zebrafish kidney. The counts were then adjusted using Poissonian counting statistics to yield the expected number of molecules as described in.

Data was normalized by median normalization to a minimum number of 1000 transcripts and genes expressing at least three transcripts in at least two cells were retained for zebrafish WKM. Pancreatic data was median normalized to 4000 transcripts and only genes expressing 5 transcripts in at least 3 cells were retained for downstream analysis. We then computed the Pearson’s distance (1 - *p*) between cells. To cluster cells, we used a method previously published in ref. 6. Briefly, we used hierarchical clustering (‘hclust’ R function with ‘ward.D2’ method) to cluster cells. To identify the number of clusters, we used ‘cutreeDynamic’ along with the ‘hybrid’ method which allows the user to specify a ‘deepSplit’ parameter controlling the sensitivity of clustering. We evaluated 100 subsamples of our data by randomly selecting 90% of the genes in the dataset, specifying the ‘deepSplit’ parameter as an integer from 0 to 4 and evaluating the average silhouette width of the number of clusters. This procedure resulted in identifying the correct cell types for both data sets of the zebrafish WKM data and the pancreatic data.

While evaluating the results of our enrichment experiments, we clustered all data together to ensure maximum confidence in resulting purity estimates. For zebrafish, this involved clustering both training datasets and enrichment experiments (WKM 1-15) resulting in 15984 cells in all. For the pancreas data, clustering both training data sets and data from four donors resulted in a total of 10176 cells.

Differentially expressed genes between two subgroups of cells were identified similar to a previously published method^7^. Briefly, we started by modeling the background expected transcript count variability. We then identified genes in each subgroup that were variably expressed by representing gene expression of each gene as a negative binomial distribution. We then computed Benjamini-Hochberg corrected *p*-values for the observed difference in transcript counts between the two subgroups as described earlier^8^ and identified differentially expressed genes (adjusted *p*-value < 0.01). Such genes were then used to annotate specific cell types within each model system based on known published literature.

For the zebrafish WKM data, we selected the topmost ten genes for each cell type ordered by their log fold change in expression when comparing the gene’s expression in a specific cell cluster compared to other cell clusters taken together (Supplementary figure 1c). Some known marker genes, especially for HSPCs and lymphocytes do not make the top ten list. We manually added them to our list of differentially expressed genes. We then used hierarchical clustering to cluster genes in seven clusters (one for each cell type). We found that our manually added genes, namely, *meis1b, myb* (denoting HSPCs^9^) and *pax5, cd79b* (denoting lymphocytes^9^) clustered in the appropriate clusters and do not show expression elsewhere (Supplementary figure 1c).

### Gate prediction methodology

The goal of GateID is to predict gates towards sorting a desired cell type from a mixture of multiple cell types. In other words, we want to purify a specific cell type to maximum purity while sorting a sufficient fraction of the desired cells. Recent advances in flow cytometry allow users to index sort, which is to save and associate flow cytometry readouts pertinent to each sorted cell. After performing single-cell mRNA sequencing, one can then merge this information with the cell type annotation (Fig. 1 – Step 1) for each cell. Such a merged data set forms the starting point for GateID, and we refer to it as training data.

We treat gate prediction as an optimization problem, wherein predicted gates allow a minimal number of undesired cells while maximizing the number of desired cells. The algorithm takes as input a matrix with FACS measurements and cell type annotation for each cell. It requires the desired cell type and the minimum yield to be input by the user. Yield is defined as a percentage of desired cells (of the total number of desired cells) that are predicted to pass through the gates. GateID first predicts a gate for each pair of flow cytometer channels, comprising scatter and fluorescence channels, where each gate is represented as a polygon with four vertices. The starting gate is computed by setting its vertices to represent the 2nd and 98th percentile in each of the *x* and *y*-axis and functions as the starting point for the optimization algorithm. We use a two-step optimization as follows for the prediction of a gate -

1. The first step finds a gate that contains at least the user-specified minimum yield for desired cells while minimizing the number of undesired cells in the gate. Fitness of each solution is thus defined by the number of undesired cells in the gate. The highest fitness is the complete absence of undesired cells within the gate. The requirement of minimum yield is enforced by assigning the worst fitness (equivalent to the total number of undesired cells in the data set) to a solution not adhering to this constraint.
2. The second step takes as input the solution (gate) of the first step and tries to maximize the yield while disallowing an increase in undesired cells. Fitness in this step is thus defined as the number of desired cells within the gate. Best fitness is achieved when all desired cells are sorted by the gate. The requirement of maximum number of undesired cells is enforced by assigning the worst fitness of zero yield to a solution not adhering to the constraint.

By default, each step is run for 20000 iterations. While evaluating fitness at each iteration, we only allow solutions involving convex polygons thereby dismissing non-convex shapes that may result in over-fitting on the training data.

Once gates for each pair of FACS channels are predicted, gate combinations can then be evaluated in logical conjunction (AND combination) such as all combinations of two gates, all combinations of three gates or a higher order. For example, many of the experiments in this study were carried out on BD FACSJazz^TM^, which records cytometry readouts in twelve channels, six scatter and six fluorescence channels. There are thus *C(12,2)* = 66 channel pairs and 66 gates. 66 gates can be further combined to yield 2.145 pairwise gate combinations *(C(66,2))* evaluated in an AND configuration, meaning a cell has to pass through both gates to be sorted.

A possibly better strategy could be to optimize a pair of gates in AND combination together, because optimization together may allow an increase in yield while reducing impurity in a coordinated fashion. One can thus optimize all 2.145 combination of pairwise gates together. In all examples we tested, two gates were enough to achieve high purity. These include stained samples of HSPCs, lymphocytes, eosinophils and monocytes. While optimizing all 2145 pairwise gates is possible, the number quickly explodes thereafter to 45760 (combinations of 3 gates) and 720720 for 4 gate combinations and thus may become intractable.

This leads us to the third intuitive approach - that of recursive gating: once gates for each combination of FACS channels are predicted (66 gates in this study), the best gate in terms of purity is selected. This gate is paired with each other gate and re-optimized together. This process is repeated until 100% purity is reached, no overall improvement is observed in the subsequent iteration or if the number of gates exceeds a user-defined preset limit. This method is added in the algorithm but hasn’t been tested experimentally.

Even if there are differences in methods mentioned above, different approaches predict gates that are comparable in yield and purity, demonstrated by experiments enriching eosinophils from the unstained sample (Fig 2), wherein the first method was used versus experiments enriching eosinophils from the stained sample, where each pair of gates was optimized together (Supplementary fig. 3c-f). Both experiments predicted similar purities for enrichment of eosinophils. Gates for pancreatic cell types were predicted by using the first method of optimizing gates separately, which predicted and achieved high purities experimentally (Fig. 4, Supplementary Fig. 9-10).

As stated above, the objective function of the optimization procedure is to predict gates that allow a minimal number of undesired cells while maximizing the number of desired cells. This presents a discrete problem for optimization. In addition, scRNA-seq along with flow cytometry results in a limited number of cells, wherein the complete variance of each cell type population may not be captured sufficiently, especially for rarer cell populations. To address these problems, we chose a derivative-free, fast and robust optimization algorithm called MA-LS- Chains, which combines an evolutionary algorithm along with a local search and is available as an R package (Rmalschains^10^). Such algorithms are known to converge faster and more reliably without being trapped in local optima (references within ^10^). While theoretically any robust global optimization algorithm may suffice, a comparison with other algorithms (Supplementary Fig. 12, and see below) shows that MA-LS-Chains is both fast and optimizes to the best purity. This is not surprising in the light of the “no free lunch” theorems, which state that certain optimization algorithms may do better than others for a certain kind of problem^11^.

The procedure above states in brief how gates are predicted. However, every sorted biological sample is different owing to multiple sources of variability. For example, variability is introduced during tissue isolation and subsequent sorting. An added layer of variability springs from fluctuating proportion of each cell type per isolation and variability in the statistical properties for each cell type in FACS space. For instance, the inconsistency in the proportion of each cell type can be readily observed by comparing the unenriched barplots in Supplementary figures 3e-f and 6c-e for the zebrafish, and Supplementary fig. 10f-g for human pancreas. Such inconsistency is further exacerbated by an overall shift in the distribution of all points demonstrated in Supplementary Fig. 2a (WKM1-3). For example, the distribution of side scatter height changes from a maximum of ~500 (WKM1) to ~100 (WKM2 and WKM3). Such variability requires that GateID predicted gates also change with respect to the current experiment in real time (Fig 1 – Step 3f).

The first approach, normalization method 1, is to deal with such variability by standardizing the values for the vertices of gates to the unenriched population of cells of the training data set using z-normalization. During FACS enrichment, one can analyze sufficient events (~10000 events) and use the mean and the standard deviation of the population of the current sort to normalize gates using the reverse of z-normalization procedure. Another method for gate normalization is elaborate and requires machine learning, and we refer to it as normalization method 2. Briefly, one first trains a machine learning classifier to classify the desired cell type based on the training data. The current sort, however, could have a different overall distribution of points, different cell type proportion therefore changing the statistical variance in different dimensions and is known to create a problem for classifiers^12^. Thus, data from the current sort needs to be normalized to the target distribution of the training data for each FACS channel. To do this, one can use methods such as non-linear qspline normalization^13^ used to compare different microarray chips to each other. Once the new data is normalized in this fashion, we classify the cells therein as desired and undesired cells using the trained classifier. We next z-normalize predicted gates to the desired cells from our training data and renormalize them to the predicted desired cells in the new data from the current sort. We again use z-normalization, but instead of normalizing the gates to the complete dataset, we normalize them using only the predicted desired population. This approach accounts for high variability in cell-type proportions from experiment to experiment, as opposed to the reverse z-normalization strategy on the complete set of points, which accounts for overall variability in the distribution of the whole data set. We note that one can also use normalization method 2 without the use of qspline normalization but with machine learning included to train and then predict desired cells from the current FACS enrichment experiment.

### Gate prediction for zebrafish WKM and Human pancreatic alpha and beta cells

To predict gates for eosinophils from the unstained zebrafish WKM, we used GateID to optimize gates on each of the pairs of FACS channels (66 gates) and then computed the best combination of two gates in an AND combination (Supplementary table 1). Gates were normalized using the normalization method 1 for each of the eosinophil sorts from the unstained WKM. For experiments concerning hematopoietic cell types in the stained WKM on BD FACSJazz™ (HSPCs, lymphocytes, monocytes and eosinophils), we optimized all 2145 gate combinations together in a pairwise fashion. For experiments on BD FACSInflux™, we optimized 61425 gate combinations together in a pairwise fashion. As one can observe from the eosinophil enrichment, both methods yielded experimentally similar results (Fig 2d, 3b). Gates were normalized for each sort using normalization method 2.

For alpha and beta cells from the human islets of Langerhans, we optimized gates for each of the pairs of FACS channels (66 gates) and computed the best combination of gates in AND configuration. Gates for alpha cell and beta cells predicted from the smaller training data (d30, Supplementary fig. 9) were normalized using the mean normalization method 1 relying on the whole population of cells, as were the beta cell gates for the second donor, based on the second training dataset (Supplementary fig. 10a-e). To compare normalization methods, beta cells from the third donor were normalized using both normalization method 1 and 2 that yielded similar results. We displayed the results from method 2 in Supplementary Fig. 10.

### Comparison of different optimization algorithms

Different optimization algorithms may perform variably for different optimization tasks. To check if our choice of using MA-LS-Chains was indeed the best, we evaluated eight different optimization algorithms (Supplementary Fig. 12). These were controlled random search (CRS, R package: nloptr^14,15^), continuous genetic algorithm (GA, R package: GA^16^), MA-LS-Chains (R package: Rmalschains^10^), bounded Hooke-Jeeves (HJK, R package: dfoptim^17^), bounded Nelder-Mead (NMK, R package: dfoptim^17^), simulated annealing (SA, R package: GenSA^18^), DEoptim (R package: RcppDE^19^), bound optimization with quadratic approximation (BOQA, R package: nloptr^20^). We randomly chose two gates to optimize together using the stained WKM and HSPCs as the desired cells. For each optimization algorithm, we optimized those gates for maximum purity with at least a 20% yield. We repeated this process 100 times while choosing two random gates to optimize every iteration and recorded the purity of each optimization algorithm.

### Computational generation of inflated dataset(s) for understanding the size of training data

It is important to understand the size of training dataset required for the generation of robust gates. Here, we wish to make a distinction between the feasibility of designing a gate and its efficacy during a real experiment. GateID can design gates with little number of cells, as in the case of eosinophils from the unstained training data 1 for the zebrafish WKM. In this particular case, there are 48 eosinophils in the training dataset (3.8% of total cell composition). Eosinophils are relatively distinct in FACS space allowing GateID to predict gates with high-predicted purity. However, an enrichment experiment involves many more cells with higher variance in their FACS readouts that may not always be represented in the training dataset. If this is the case, GateID cannot take into account FACS profiles for possible contaminating cell types that are not visible to the algorithm while predicting gates. Thus in practice, GateID predicted gates may not perform as predicted using a smaller dataset. We believe this is the reason behind the fact that the predicted beta cell gates designed on the first limited training dataset (664 cells) of the pancreas did not do well in an actual experiment (Fig. 4b, Supp. fig 9d- e, donor 1 – 78.3% purity). To further check if our hypothesis was true, we did the following computational experiment. We used our limited training dataset from the pancreas (training dataset 1, 664 cells) to generate two larger datasets using truncated multivariate sampling. Briefly, we sampled random instances from a normal distribution parameterized by the mean and variance of each cell cluster in the dataset, for each FACS channel. We used this method to increase the size of our dataset twofold and then threefold. We took care that the random deviates resided within the bounds of zero and a maximum of the particular FACS channel, similar to data from a FACS experiment. This method also ensured that the artificial datasets would have identical proportion of cell clusters in comparison to the training data. We then used GateID to design gates on both these artificial datasets.

To evaluate the performance of the gates generated above, it is important to take into account that contaminating cells may be higher in number in another experiment. We therefore listed the most common cell type(s) that contaminates beta cell gates (alpha cells and delta cells for our dataset) and increased its proportion in stepwise fashion while generating a dataset of a larger size. Here, we used a size of 20000 cells, which is in the same range as an actual experiment involving sorting of live cells. We then evaluated the gates predicted by GateID on our actual training dataset 1 (664 cells) and two artificial training datasets on this test dataset. We repeated this evaluation test 50 times for each of the three sets of gates. We observed that gates designed on a smaller dataset (1x) fare poorly in comparison to gates designed on a larger dataset (2x and 3x) (**Supplementary Fig. 11b**). Specifically, increasing the training dataset two fold to 1328 cells ensures higher mean purity even in the case of twice the amount of contaminating cells and may ensure higher robustness to fluctuations in cell proportions.

Truncated multivariate sampling was carried out using package ‘tmvtnorm’ in R.

## Acknowledgements

This work was supported by a European Research Council Advanced grant (ERC-AdG 742225-IntScOmics) and Nederlandse Organisatie voor Wetenschappelijk Onderzoek (NWO) TOP award (NWO-CW 714.016.001). This work is part of the Oncode Institute, which is partly financed by the Dutch Cancer Society. We especially thank Josi Peterson-Maduro, Lennart Kester and all the other members of the A.v.O. laboratory for discussions and input. In addition, we thank the Hubrecht Sorting Facility, and the Utrecht Sequencing Facility, subsidized by the University Medical Center Utrecht, Hubrecht Institute and Utrecht University.

## Author contributions

A.v.O and A.B conceived and designed the project. A.B developed the GateID algorithm. A.B, C.S.B, M.J.M and A.v.O further refined the algorithm. A.B performed the gate design and normalization for BD FACSJazz™ WKM and pancreas experiments. C.S.B performed the gate design and normalization for BD FACSInflux™ WKM experiments. R.v.d.L operated both FACS machines used in this study. C.S.B performed zebrafish WKM scRNA-Seq experiments. A.B. and C.S.B analyzed the zebrafish WKM scRNA-Seq data. M.J.M performed human pancreas scRNA-Seq experiments. M.J.M analyzed the human pancreas scRNA-Seq data. G.D and E.d.K provided human pancreatic tissue. All authors discussed and interpreted the results. C.S.B, A.B, M.J.M and A.v.O wrote the manuscript. C.S.B, A.B and M.J.M contributed equally to this work.

## Data Availability

The accession numbers for the scRNA-Seq datasets reported in this study are available on GEO: GSE112438.

