## Supplementary figures and tables for "Cell type purification by single-cell transcriptome-trained sorting"

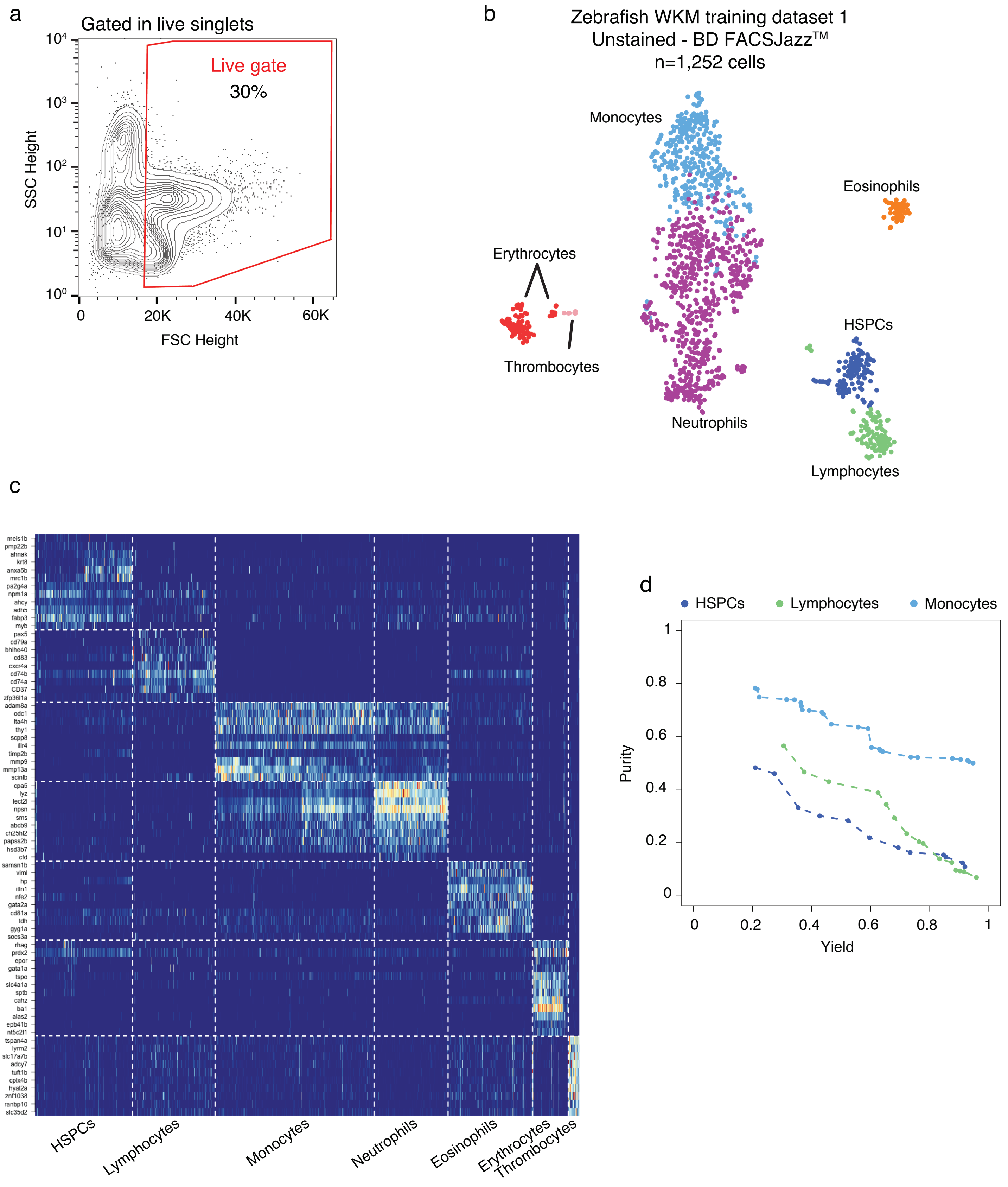


Baron et al, Supplementary Figure 1

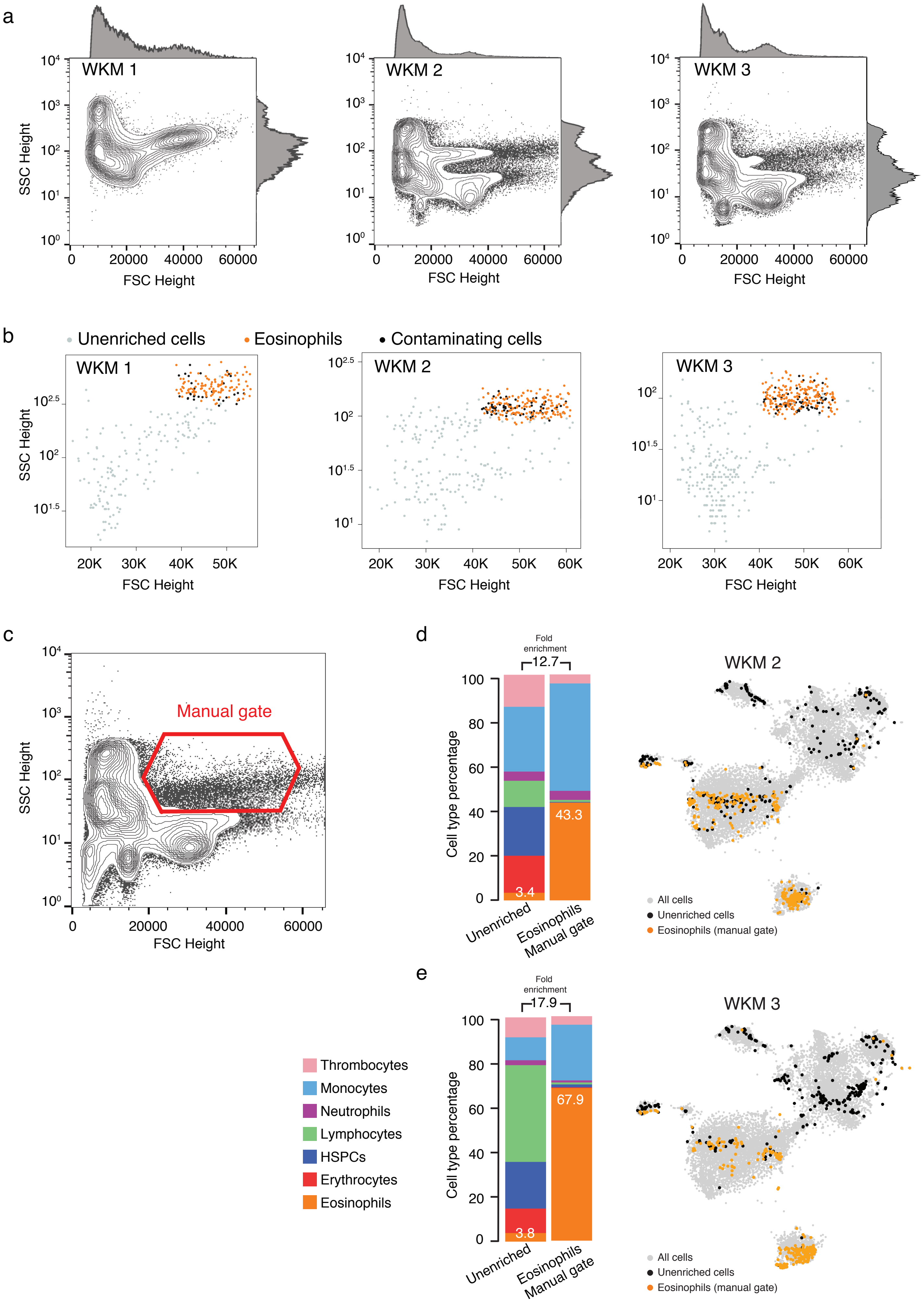


Baron et al, Supplementary Figure 2

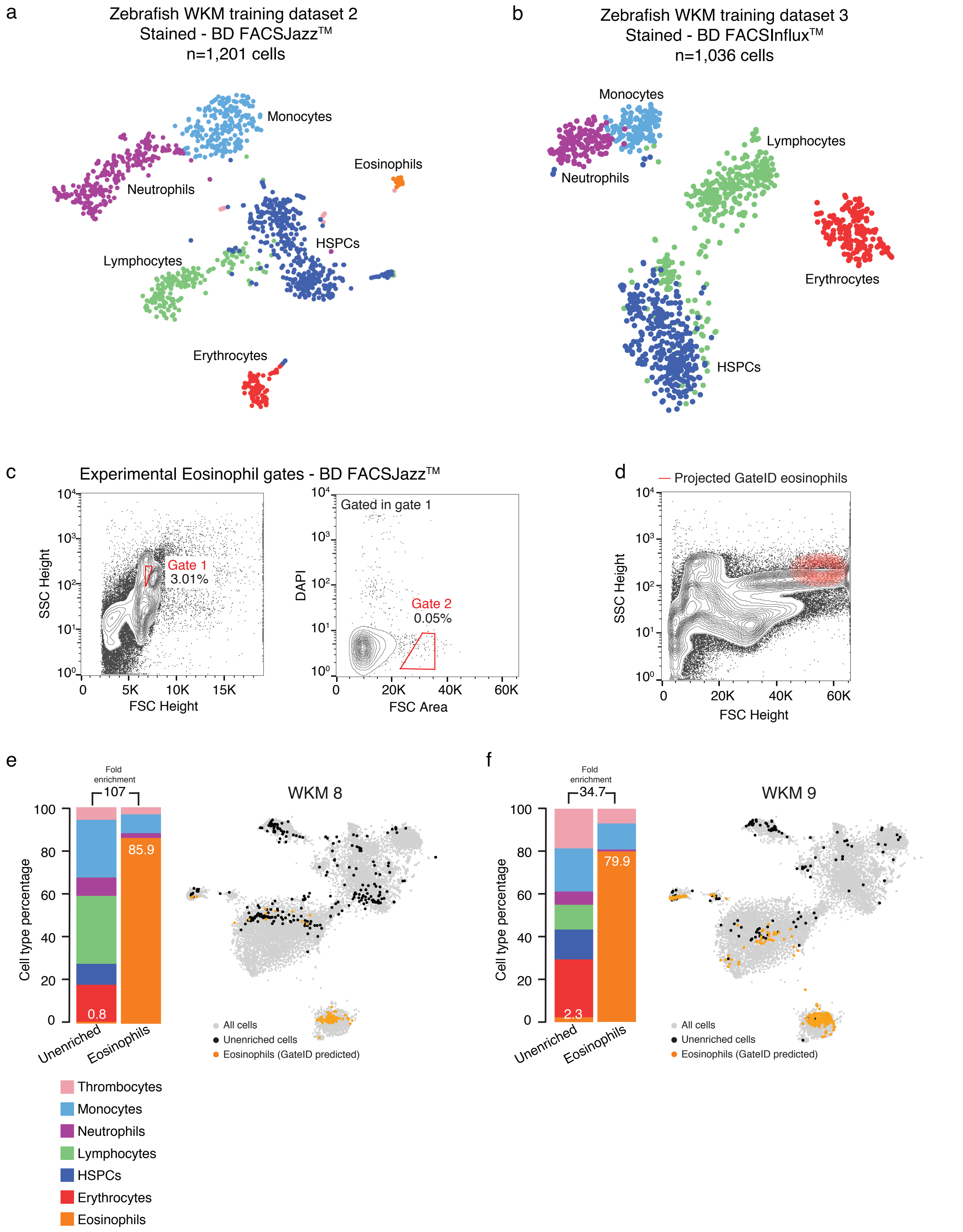


Baron et al, Supplementary Figure 3

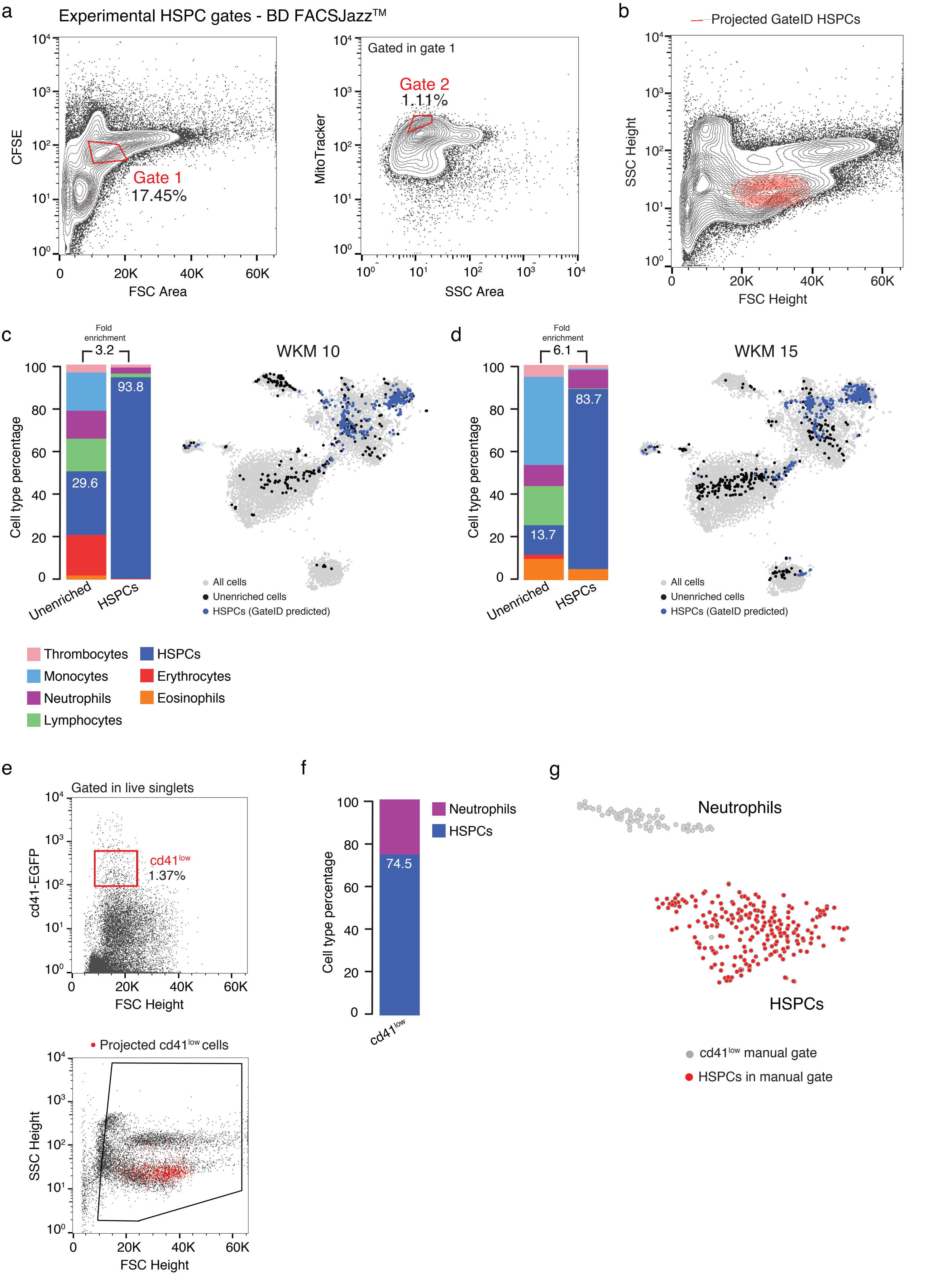


Baron et al, Supplementary Figure 4

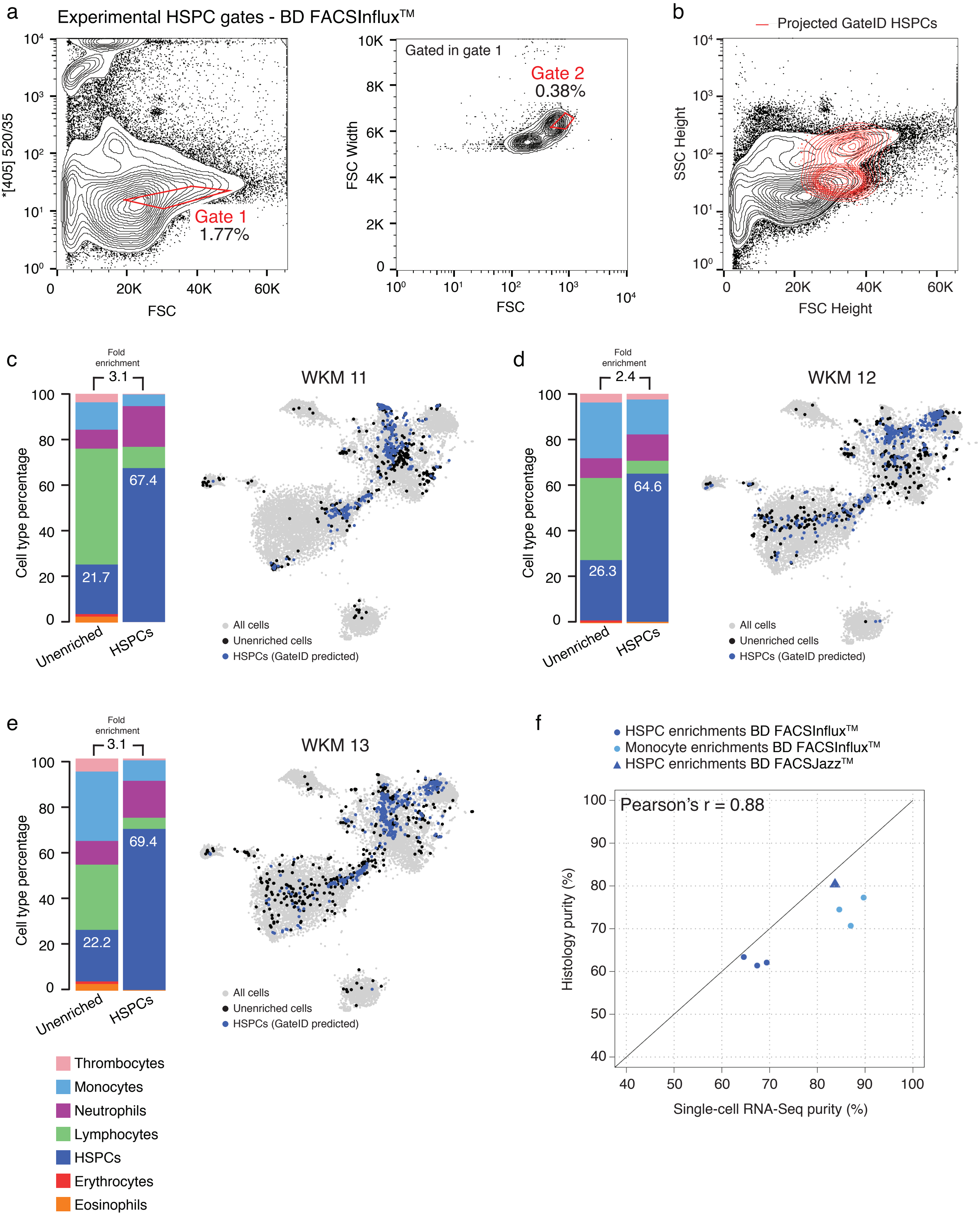


Baron et al, Supplementary Figure 5

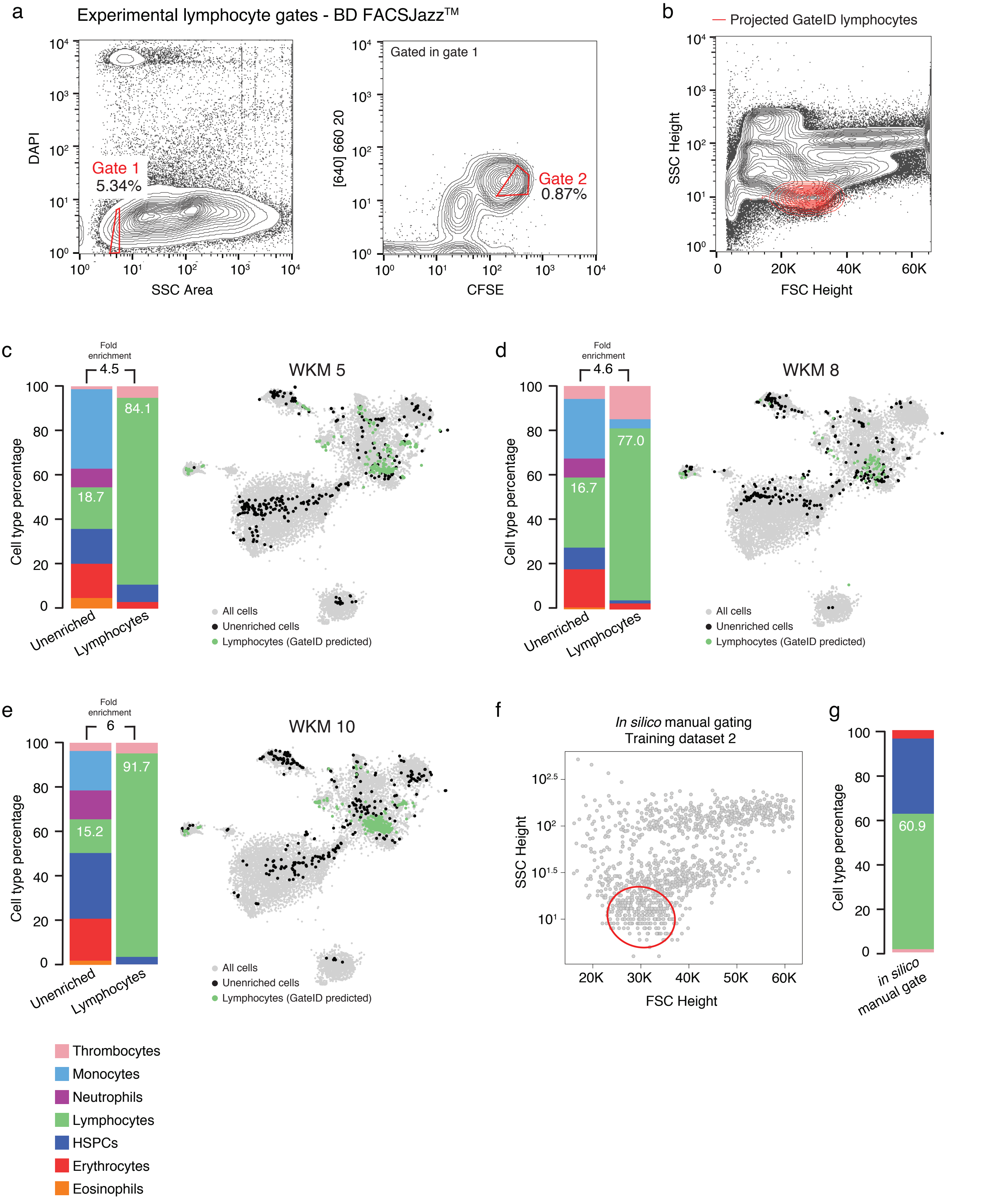


Baron et al, Supplementary Figure 6

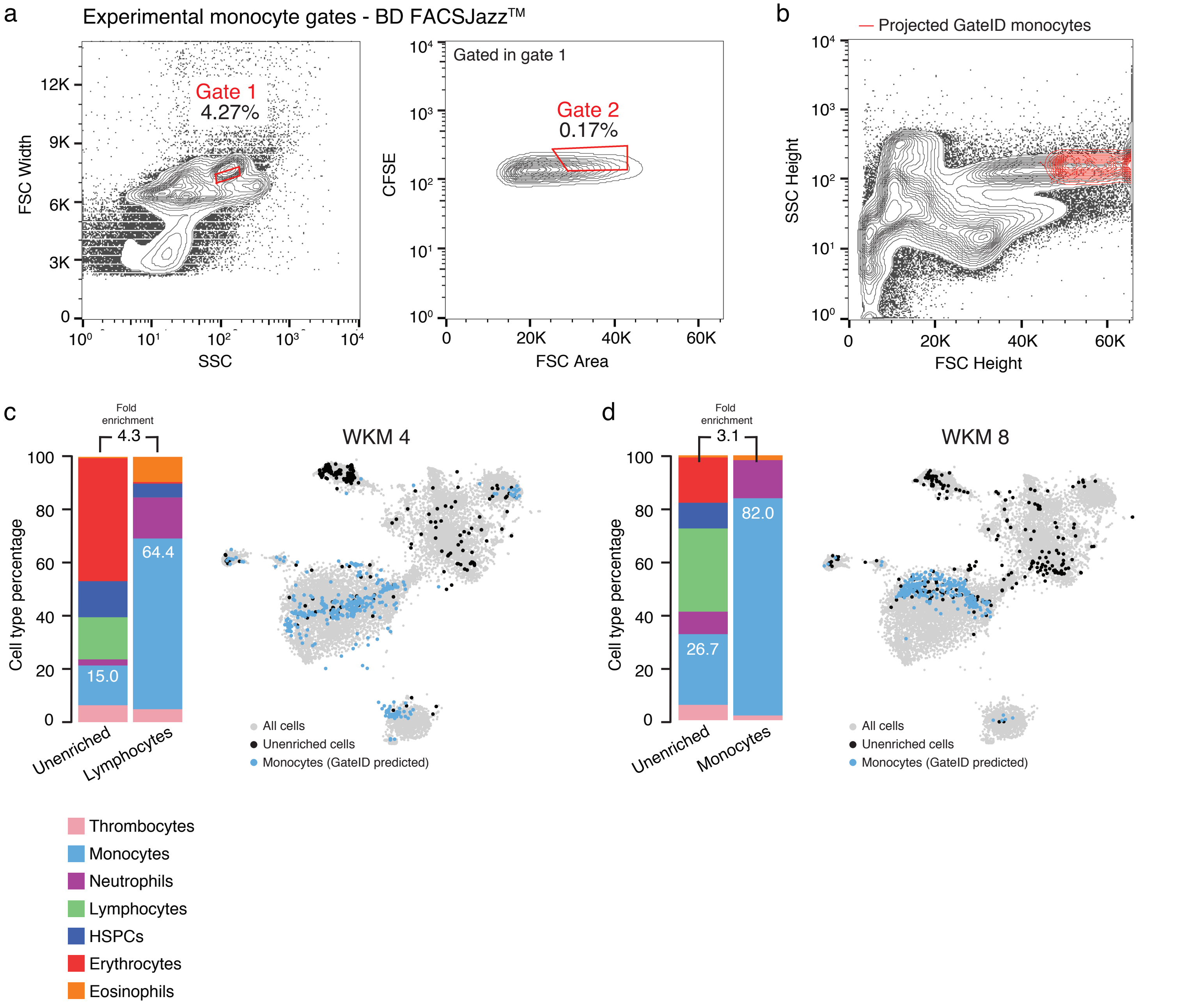


Baron et al, Supplementary Figure 7

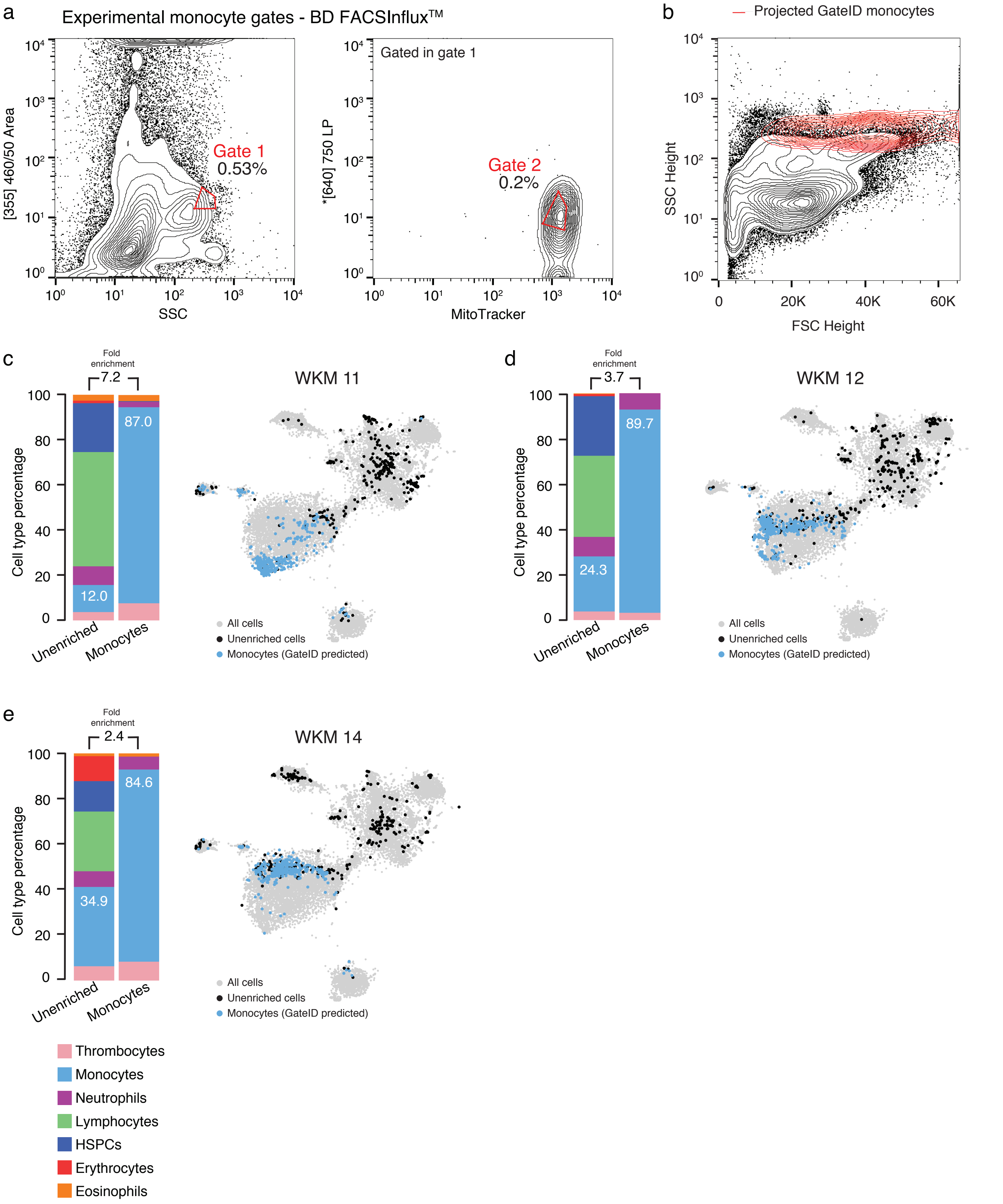
a
Experimental monocyte gates - BD FACSInflux™
[355] 460/50 Area
SSC
Gate 1
0.53%
Gated in gate 1
*[640] 750 LP
MitoTracker
Gate 2
0.2%
b
Projected GateID monocytes
SSC Height
FSC Height
0
20K
40K
60K
c
Fold enrichment
7.2
Cell type percentage
87.0
12.0
Unenriched
Monocytes
WKM 11
All cells
Unenriched cells
Monocytes (GateID predicted)
d
Fold enrichment
3.7
Cell type percentage
89.7
24.3
Unenriched
Monocytes
WKM 12
All cells
Unenriched cells
Monocytes (GateID predicted)
e
Fold enrichment
2.4
Cell type percentage
84.6
34.9
Unenriched
Monocytes
WKM 14
All cells
Unenriched cells
Monocytes (GateID predicted)
Thrombocytes
Monocytes
Neutrophils
Lymphocytes
HSPCs
Erythrocytes
Eosinophils

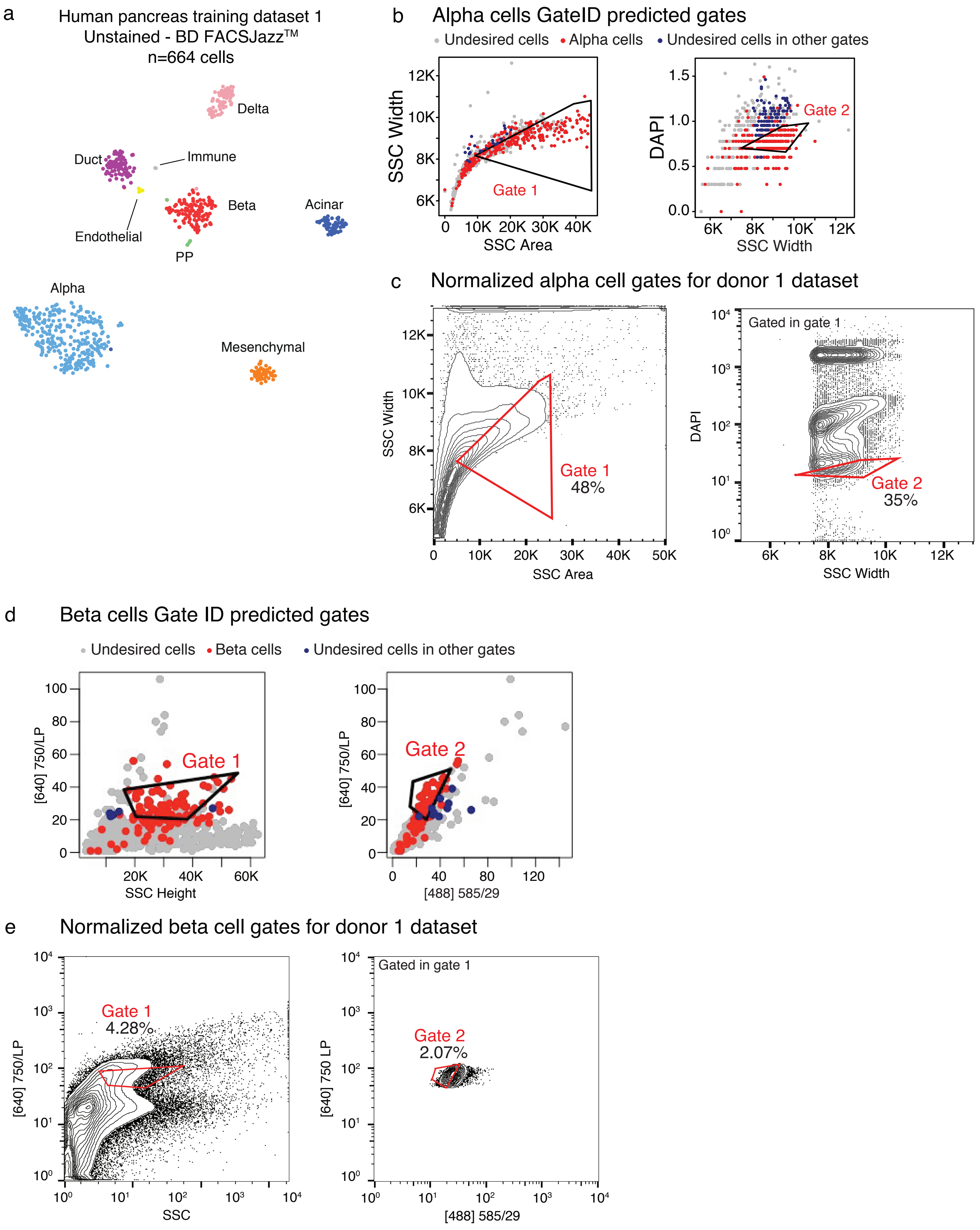


Baron et al, Supplementary Figure 9

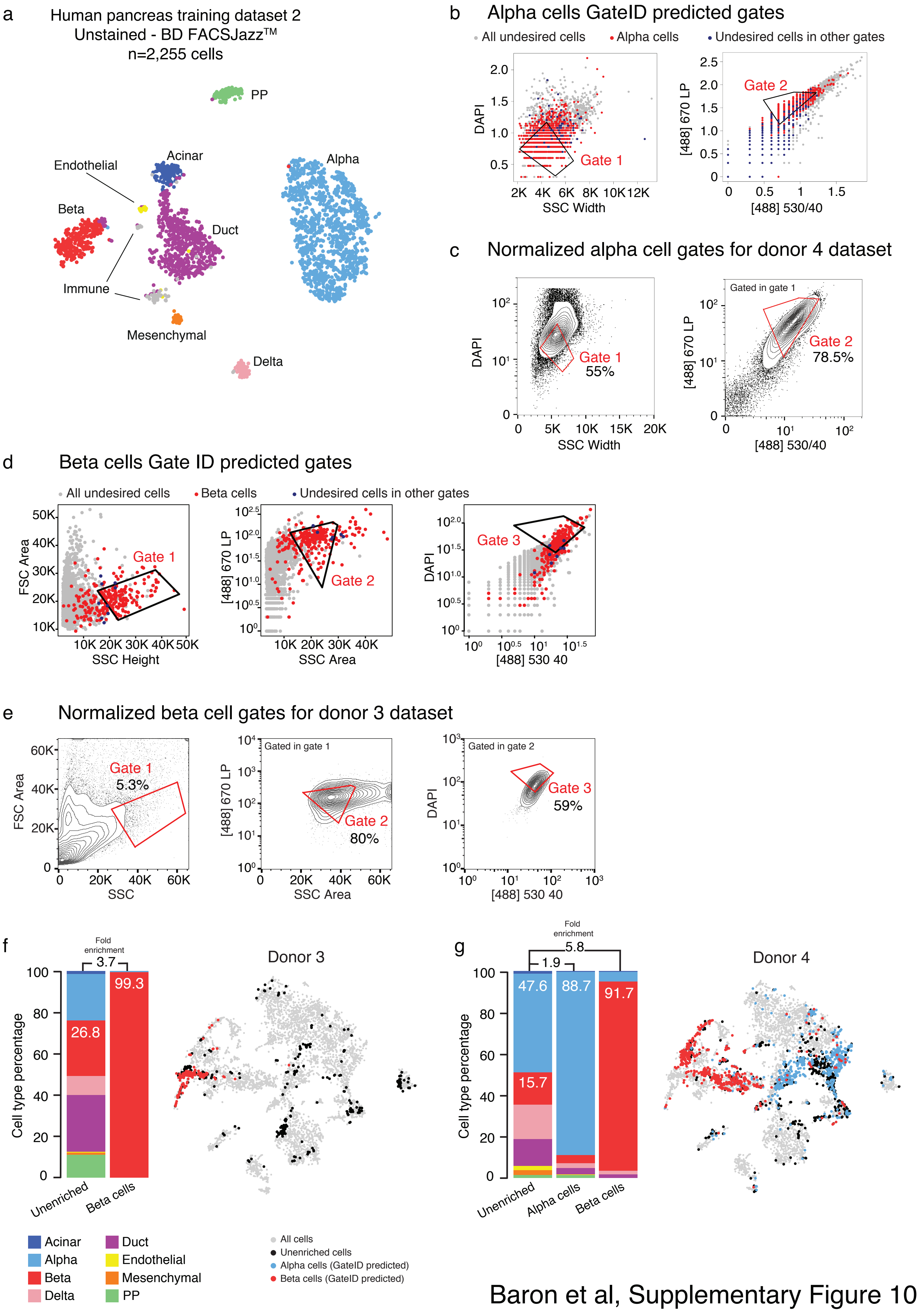


Baron et al, Supplementary Figure 10

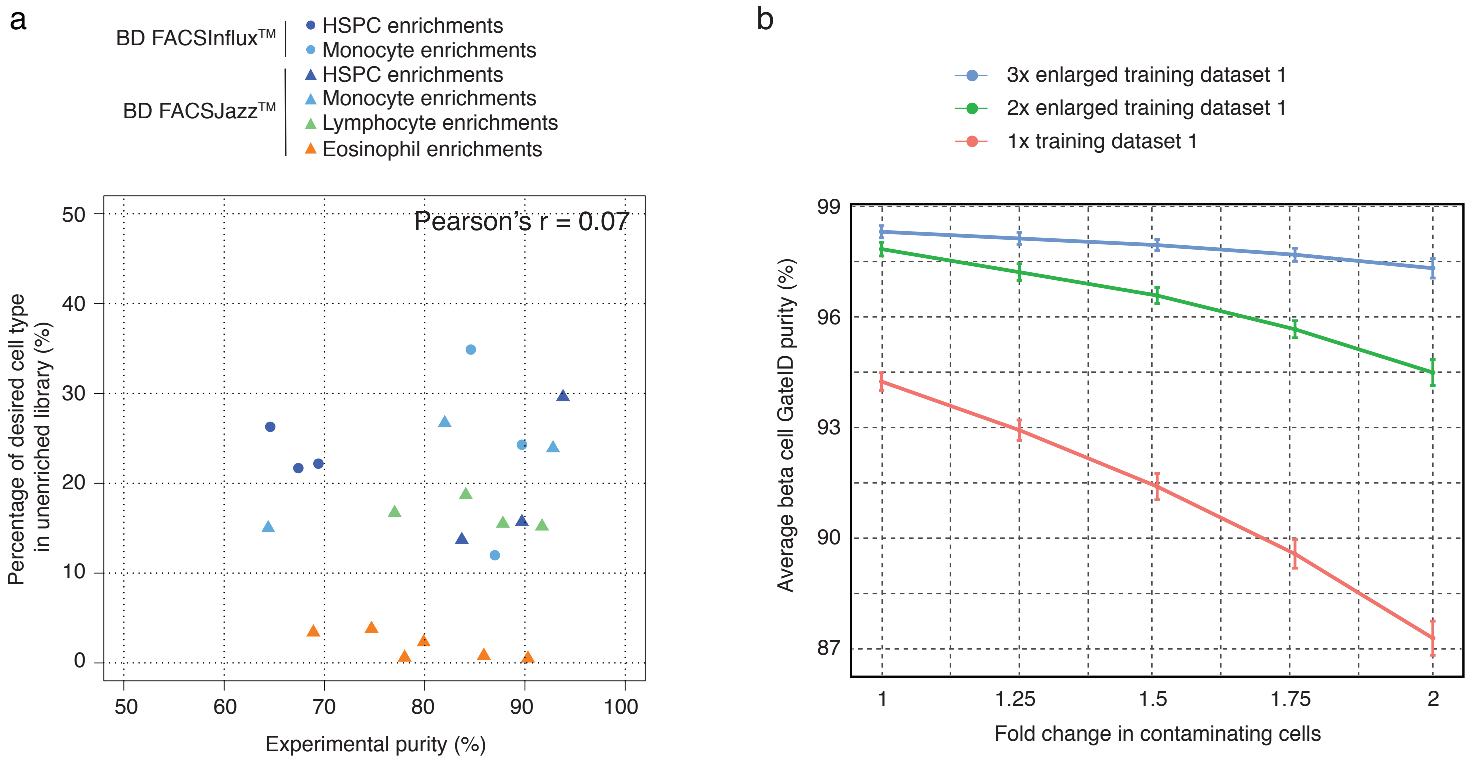


Baron et al, Supplementary Figure 11

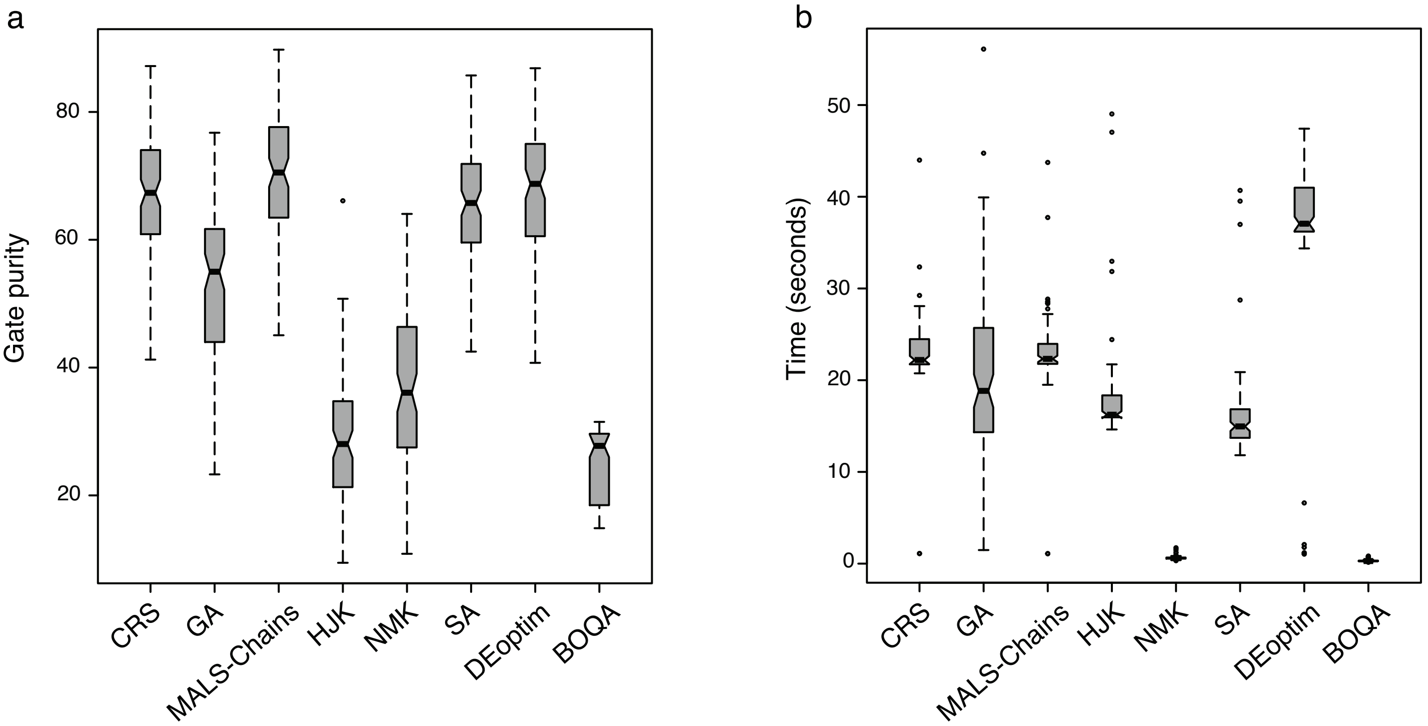


Baron et al, Supplementary Figure 12

| Experiment label | Cell type | Training dataset | FACS machine | Predicted yield (%) | Predicted purity (%) | Sorted fraction in unenriched (%) | Experimental purity (%) |
|---|---|---|---|---|---|---|---|
| **Zebrafish WKM enrichments** | | | | | | | |
| WKM 1 | Eosinophils | 1 (unstained) | BD FACSJazz™ | 46.9 | 79.3 | 0.6 | 78 |
| WKM 2 | | | | | | 3.4 | 68.9 |
| WKM 3 | | | | | | 3.8 | 74.7 |
| WKM10 | HSPCs | 2 (stained) | | 20 | 90.5 | 29.6 | 93.8 |
| WKM5 | | | | | | 15.7 | 89.7 |
| WKM15 | | | | | | 13.7 | 83.7 |
| WKM10 | Lymphocytes | | | 20 | 97.5 | 15.2 | 91.7 |
| WKM5 | | | | | | 18.7 | 89.7 |
| WKM8 | | | | | | 16.7 | 77 |
| WKM6 | | | | | | 15.5 | 87.8 |
| WKM 8 | Monocytes | | | 20.5 | 98 | 26.7 | 82 |
| WKM 4 | | | | | | 15 | 64.4 |
| WKM 7 | | | | | | 23.9 | 92.8 |
| WKM 8 | Eosinophils | | | 23 | 100 | 0.8 | 85.9 |
| WKM 9 | | | | | | 2.3 | 79.9 |
| WKM 4 | | | | | | 0.5 | 90.3 |
| WKM11 | Monocytes | 3 (stained) | BD FACSInflux™ | 40 | 100 | 12 | 87 |
| WKM12 | | | | | | 24.3 | 89.7 |
| WKM14 | | | | | | 34.9 | 84.6 |
| WKM11 | HSPCs | | | 30 | 98.6 | 21.7 | 67.4 |
| WKM12 | | | | | | 26.3 | 64.6 |
| WKM13 | | | | | | 22.2 | 69.4 |
| **Human pancreas enrichments** | | | | | | | |
| Donor 1 | Alpha cells | 1 (unstained) | BD FACSJazz™ | 42.6 | 100 | 44.4 | 97.2 |
| | Beta cells | | | 51.7 | 100 | 6.1 | 78.3 |
| Donor 2 | Beta cells | 2 unstained | | 25.8 | 98.4 | 25 | 94.8 |
| Donor 3 | Beta cells | | | 25.8 | 98.4 | 26.8 | 99.3 |
| Donor 4 | Alpha cells | | | 51.2 | 97.12 | 47.6 | 88.7 |
| | Beta cells | | | 25.8 | 98.4 | 15.7 | 91.7 |

Supplementary table 1

**Supplementary Figure 1. Generation of zebrafish WKM unstained training dataset**

**(a)** FSC Height and SSC Height contour plot of sorted live WKM cells to generate WKM training dataset 1. Events represented are live singlets from the total WKM population. Majority of erythrocytes were excluded by excluding events with low FSC-Height. **(b)** t-SNE map of zebrafish WKM training dataset 1 generated on BD FACSJazz$^{TM}$. Single cells are colored based on cell type. **(c)** Heat map showing marker genes for all hematopoietic cell types identified in the WKM full dataset. **(d)** Curves showing trade-off between yield and purity of GateID solutions for HSPCS, lymphocytes and monocytes for the unstained training dataset 1. All gates for a given cell type with lower purity or yield are internal to these curves and are not shown.

**Supplementary Figure 2. Eosinophil enrichments with unstained WKM cells on BD FACSJazz$^{TM}$**

**(a)** FSC Height and SSC Height contour plots of all WKM cells analyzed for eosinophils enrichment experiments WKM 1 to 3. Histograms on each plot show population density is FSC and SSC Height channels. **(b)** Plots showing sorted unenriched and GateID enriched cells for eosinophil experiments WKM 1 to 3 in FSC and SSC Height. Grey points are cells from the unenriched library and colored points are cells from the GateID enriched library. Sorted eosinophils in the GateID enriched library are highlighted in orange and sorted non-eosinophil contaminating cells in the GateID enriched library are represented in black **(c)** FSC Height and SSC Height contour plot of all WKM cells for WKM 2. The eosinophil manual gate used in WKM 2 experiment is represented in red (representative for WKM 2 and 3 manual enrichment experiments). **(d-e)** Barplots and t-SNE maps showing the outcome of eosinophil enrichments using manual gating for two independent experiments (WKM 1 and 2) on BD FACSJazz$^{TM}$. In the barplots, numbers in the bars indicate the percentage of eosinophils in the corresponding library and numbers above the bars indicate the cell type fold enrichment between unenriched and manually enriched library. On the t-SNE maps, grey points represent all cells from the WKM dataset. For each experiment, black dots are single cells in the unenriched library for a given experiment, while colored dots are single cells in the manually enriched library for the same experiment.

**Supplementary Figure 3. Eosinophil enrichments with stained WKM cells on BD FACSJazz$^{TM}$**

**(a)** t-SNE map of zebrafish stained WKM training dataset 2 generated on BD FACSJazz$^{TM}$. Single cells are colored based on cell type. **(b)** t-SNE map of zebrafish stained WKM training dataset 3 generated on BD FACSInflux$^{TM}$. Single cells are colored based on cell type. **(c)** Contour plots of stained WKM cells showing normalized sorting gates for eosinophils for WKM 8 experiment (representative example for WKM 4, WKM 8 and WKM 9 eosinophil enrichments) on BD FACSJazz$^{TM}$. Sorted cells passed through gate 1 and gate 2. Percentages of events within each gate are indicated. **(d)** Projection of the sorted GateID eosinophils in WKM 8 (representative example for WKM 4, 8 and 9 eosinophil enrichment experiments) in FSC Height vs. SSC Height. **(e-f)** Barplots and t-SNE maps showing the outcome of eosinophil enrichments for **(e)** WKM 8 and **(f)** WKM 9 on BD FACSJazz™. In the barplots, numbers in the bars indicate the percentage of eosinophils in the corresponding library and numbers above the bars indicate the eosinophil fold enrichment between unenriched and GateID enriched library. On the t-SNE maps, grey points represent all cells from the WKM dataset. For each experiment, black dots are single cells in the unenriched library for a given experiment, while colored dots are single cells in the GateID enriched library for the same experiment.

**Supplementary Figure 4. HSPC enrichments with stained WKM cells on BD FACSJazz™**
**(a)** Contour plots of stained WKM cells showing experimental sorting gates for HSPC for the WKM 10 experiment (representative example for WKM 5, 10 and 15 HSPC enrichment experiments) on BD FACSJazz™. Sorted cells passed through gate 1 and gate 2. Percentages of events within each gate are indicated. **(b)** Projection of the sorted GateID HSPCs for WKM 10 in FSC Height vs. SSC Height (representative example for WKM 5, 10 and 15 HSPC enrichment experiments). **(c-d)** Barplots and t-SNE maps showing the outcome of HSPC enrichments for **(c)** WKM 10 and **(d)** WKM 15 on BD FACSJazz™. In the barplots, numbers in the bars indicate the percentage of HSPCs in the corresponding library and numbers above the bars indicate the HSPC fold enrichment between unenriched and GateID enriched library. On the t-SNE maps, grey points represent all cells from the WKM dataset. For each experiment, black dots are single cells in the unenriched library for a given experiment, while colored dots are single cells in the GateID enriched library for the same experiment. **(e)** Upper panel - FSC Height vs. cd41-EGFP dot plot of live singlet WKM cells. The cd41$^{low}$ gate is represented in red. Lower panel - projection of the cd41$^{low}$ sorted cells in FSC Height vs. SSC Height. **(f)** Barplot indicating cell type percentages for sorted cd41$^{low}$ cells. Percentage in the barplot indicates HSPC percentage in the sorted library. **(g)** t-SNE map showing sorted cd41$^{low}$ cells. Non HSPCs are represented in grey and HSPCs in red.

**Supplementary Figure 5. HSPC enrichments with stained WKM cells on BD FACSInflux™**
**(a)** Contour plots of stained WKM cells showing experimental sorting gates for HSPC for the WKM 11 experiment (representative example for WKM 11, 12 and 13 HSPC enrichment experiments) on BD FACSInflux™. Sorted cells passed through gate 1 and gate 2. Percentages of events within each gate are indicated. **(b)** Projection of the sorted GateID HSPCs for WKM 10 in FSC Height vs. SSC Height (representative example for WKM 11, 12 and 13 HSPC enrichment experiments) **(c-e)** Barplots and t-SNE maps showing the outcome of HSPC enrichments for **(c)** WKM 11, **(d)** WKM 12 and **(e)** WKM 13 on BD FACSInflux™. In the barplots, numbers in the bars indicate the percentage of HSPCs in the corresponding library and numbers above the bars indicate the HSPC fold enrichment between unenriched and GateID enriched library. On the t-SNE maps, grey points represent all cells from the WKM dataset. For each experiment, black dots are single cells in the unenriched library for a given experiment, while colored dots are single cells in the GateID enriched library for the same experiment. **(f)** Scatter plot showing experimental purities of GateID predicted gates determined by scRNA-Seq (x axis) and histological analysis (y axis) for HSPCs (dark blue) and monocytes (light blue) on BD FACJazz™ (triangle) and BD FACSInflux™ (circle).

**Supplementary Figure 6. Lymphocyte enrichments with stained WKM cells on BD FACSJazz™**

**(a)** Contour plots of stained WKM cells showing experimental sorting gates for lymphocytes for WKM 10 experiment (representative example for WKM 5, 6, 8 and 10 lymphocyte enrichment experiments) on BD FACJazz™. Sorted cells passed through gate 1 and gate 2. Percentages of events within each gate are indicated. **(b)** Projection of the sorted GateID lymphocytes for WKM 10 in FSC Height vs. SSC Height (representative example for WKM 5, 6, 8 and 10 lymphocyte enrichment experiments). **(c-e)** Barplots and t-SNE maps showing the outcome of lymphocyte enrichments for **(c)** WKM 5, **(d)** WKM 8 and **(e)** WKM 10 on BD FACJazz™. In the barplots, numbers in the bars indicate the percentage of lymphocytes in the corresponding library and numbers above the bars indicate the lymphocyte fold enrichment between unenriched and GateID enriched library. On the t-SNE maps, grey points represent all cells from the WKM dataset. For each experiment, black dots are single cells in the unenriched library for a given experiment, while colored dots are single cells in the GateID enriched library for the same experiment. **(f)** Design of *in silico* reconstruction of the manual gate for lymphocyte enrichment. Cells from WKM training dataset 2 are represented in grey and manual gate is drawn in red. **(g)** Barplots indicating cell type percentages for the lymphocyte *in silico* manual gate. Percentage in the barplot indicates lymphocyte percentage in the *in silico* manual gate.

**Supplementary Figure 7. Monocyte enrichments with stained WKM cells on BD FACSJazz™**

**(a)** Contour plots of stained WKM cells showing experimental sorting gates for monocytes for WKM 8 experiment (representative example for WKM 4, 7 and 8 monocyte enrichment experiments) on BD FACJazz™. Sorted cells passed through gate 1 and gate 2. Percentages of events within each gate are indicated. **(b)** Projection of the sorted GateID monocytes for WKM 8 in FSC Height vs. SSC Height (representative example for WKM 4, 7 and 8 monocyte enrichment experiments). **(c-d)** Barplots and t-SNE maps showing the outcome of monocyte enrichments for **(c)** WKM 4 and **(d)** WKM 8 BD FACJazz™. In the barplots, numbers in the bars indicate the percentage of monocytes in the corresponding library and numbers above the bars indicate the monocyte fold enrichment between unenriched and GateID enriched library. On the t-SNE maps, grey points represent all cells from the WKM dataset. For each experiment, black dots are single cells in the unenriched library for a given experiment, while colored dots are single cells in the GateID enriched library for the same experiment.

**Supplementary Figure 8. Monocyte enrichments with stained WKM cells on BD FACSInflux™**

**(a)** Contour plots of stained WKM cells showing experimental sorting gates for monocytes for WKM 11 experiment (representative example for WKM 11, 12 and 14 monocyte enrichment experiments) on BD FACInflux™. Sorted cells passed through gate 1 and gate 2. Percentages of events within each gate are indicated. **(b)** Projection of the sorted GateID monocytes for

WKM 11 in FSC Height vs. SSC Height (representative example for WKM 11, 12 and 14 monocyte enrichment experiments). **(c-e)** Barplots and t-SNE maps showing the outcome of monocyte enrichments for **(c)** WKM 11, **(d)** WKM 12 and **(e)** WKM 14 on on BD FACInflux$^{TM}$. In the barplots, numbers in the bars indicate the percentage of monocytes in the corresponding library and numbers above the bars indicate the monocyte fold enrichment between unenriched and GateID enriched library. On the t-SNE maps, grey points represent all cells from the WKM dataset. For each experiment, black dots are single cells in the unenriched library for a given experiment, while colored dots are single cells in the GateID enriched library for the same experiment.

**Supplementary Figure 9. Gates for enrichments of alpha and beta cells from unstained pancreatic tissue on BD FACSJazz$^{TM}$**

**(a)** t-SNE map of human pancreas training dataset 1 generated on on BD FACSJazz$^{TM}$. Single cells are colored based on cell type. **(b)** GateID predicted gates to isolate alpha cells from human pancreas. Gates were predicted on training dataset 1. Red points show desired cells (alpha cells) present in training dataset and the blue points show undesired cells falling in the other gate. **(c)** Contour plots of unstained human pancreas cells showing experimental gates used to sort alpha cells from donor 1. Sorted cells passed through gate 1 and gate 2. Percentages of events within each gate are indicated. **(d)** GateID predicted gates to isolate beta cells from human pancreas. Gates were predicted on training dataset 1. Red points show desired cells (beta cells) present in training dataset and the blue points show undesired cells falling in the other gate. **(e)** Contour plots of unstained human pancreas cells showing experimental gates used to sort beta cells from donor 1. Sorted cells passed through gate 1 and gate 2. Percentages of events within each gate are indicated.

**Supplementary Figure 10. Alpha and beta cells enrichments from unstained pancreatic tissue on BD FACSJazz$^{TM}$**

**(a)** t-SNE map of human pancreas training dataset 2 generated on BD FACSJazz$^{TM}$. Single cells are colored based on cell type. **(b)** GateID predicted gates to isolate alpha cells from human pancreas. Gates were predicted on training dataset 2. Red points show desired cells (alpha cells) present in training dataset and the blue points show undesired cells falling in the other gate. **(c)** Contour plots of unstained human pancreas cells showing experimental gates used to sort alpha cells from donor 4. Sorted cells passed through gate 1 and gate 2. Percentages of events within each gate are indicated. **(d)** GateID predicted gates to isolate beta cells from human pancreas. Gates were predicted on training dataset 2. Red points show desired cells (beta cells) present in training dataset and the blue points show undesired cells falling in the other gate. **(e)** Contour plots of unstained human pancreas cells showing experimental gates used to sort beta cells from donor 3. Sorted cells passed through gate 1 and gate 2. Percentages of events within each gate are indicated. **(f-g)** Barplots and t-SNE maps showing the outcome of GateID alpha and beta cell enrichments for two independent donors on BD FACSJazz$^{TM}$. Gates were predicted on unstained training dataset 2. In the barplots, numbers in the bars indicate the percentage of alpha and beta cells in the corresponding library and numbers above the bars indicate the cell type fold enrichment between unenriched and GateID enriched library. On the t-SNE maps, grey points represent all cells from the pancreas dataset. For each experiment, black dots are single cells in the unenriched library for a given experiment, while colored dots are single cells in the GateID enriched library for the same experiment.

**Supplementary Figure 11. Determination of optimum training dataset size**
**(a)** Scatter plot showing the percentage of the desired cell type in the unenriched library versus the achieved GateID purity in the enriched library for all WKM enrichment experiments. Points are colored based on the cell type enriched (orange for eosinophils, dark blue for HSPCs, light blue for monocytes and green for lymphocytes) and shaped based on the FACS machine used for isolation (triangles for BD FACSJazz$^{TM}$ and circles for BD FACSInflux$^{TM}$). **(b)** Average beta cell purity depending on training dataset size and proportion of contaminating cells in the training dataset. The y-axis denotes the average GateID purity and its standard deviation. The x-axis represents the fold change of the proportion of the contaminating cells in the training dataset. The curves represent different datasets: 1x is the original pancreas training dataset 1 (678 cells), while 2x and 3x datasets are enlarged by two (1.356 cells) or three (2.034 cells) fold, respectively.

**Supplementary Figure 12. Comparison of various optimization algorithms**
**(a)** Purity estimate for 100 samples of gate optimization for a pair of gates using different optimization algorithms. The figure shows that MA-LS-Chains shows the best purity in comparison to 8 different optimization algorithms used here. **(b)** Time (in seconds) 100 samples of gate optimization for a pair of gates using different optimization algorithms. NMK and BOQA algorithms are fast but at the cost of substandard solution for the gate prediction problem. **(c)** Barplots indicating cell type proportions in each sequenced library (384 cells) for unenriched and GateID beta cell enriched libraries. All experiments were clustered together to call cell types. Percentages in the barplot indicate beta cell percentages in that library. Numbers above the bars indicate the beta cell fold enrichment between unenriched and GateID enriched libraries.

**Supplementary Table 1. Overview of GateID experiments**
Supplementary Table 1 shows several sorting parameters for all zebrafish WKM and human pancreas enrichment experiments. The training dataset used for the gate design is indicated as well as the fraction of desired cells in the corresponding training dataset. The GateID predicted yield and purity are indicated for each experiment. The experimentally sorted fraction of each cell type is shown next to the experimental yield and experimental purity. The experimental purity column is equivalent to the cell type percentage shown in figures 2 - 4 and the barplots in supplementary figures 1-10.
